# Self-supervised learning for a gene program-centric view of cell states

**DOI:** 10.64898/2026.03.24.713961

**Authors:** Marie Moullet, Tomoya Isobe, Amirhossein Vahidi, Carlo Leonardi, Licyel Paulas-Condori, Christopher Soelistyo, Lloyd Steele, Kevin Chi Hao Ly, Mariana Quiroga Londoño, Nicole Mende, Emily Stephenson, Deena Iskander, Simone Webb, Issac Goh, MS Vijayabaskar, Qi Liu, Daniyal J. Jafree, Hesam Asadollahzadeh, Ciro Ramírez-Suástegui, Arpit Merchant, Kenny Roberts, Benjamin Rumney, Hon Man Chan, Ash Holland, Martin Prete, David Horsfall, Daniela Basurto-Lozada, John Y. W. Lee, Elena Winheim, April Rose Foster, Roser Vilarrasa-Blasi, Rebecca Hannah, Satveer K Mahil, Catherine Smith, Anindita Roy, Irene Roberts, Elisa Laurenti, Berthold Göttgens, Roser Vento-Tormo, Muzlifah Haniffa, Nicola K. Wilson, Mo Lotfollahi

## Abstract

Single-cell omics has extended the biological interrogation of cell state from examining the expression of individual genes to unbiased profiling of tens of thousands of genes at once. However, extracting biological insights from such high-dimensional data remains challenging. To enable downstream analyses, many computational approaches compress cell state into a single latent representation. This can obscure the structure of underlying gene programs (GP), defined as coordinated sets of biologically related genes, such as signalling pathway response modules or transcription factor targets. Here, we present Tripso, a self-supervised transformer deep learning model which learns multiple GP-specific embeddings from predefined GPs, while also enabling the discovery of novel, data-driven GPs. Tripso facilitates principled comparisons across development, disease, and experimental systems. Firstly, in a dataset of human hematopoietic cells spanning prenatal development through adulthood and aging and including newly generated data, Tripso resolved age-specific GP patterns, including elevated *JAK-STAT* activity in pediatric hematopoietic cells and postnatal shifts in *IKZF1* GP activity during B cell differentiation. Secondly, leveraging Tripso GP embeddings and comparing *in vivo* to *in vitro* data, we hypothesized and experimentally validated that inhibition of the SEC61 translocon improved maintenance of hematopoietic stem cells in culture. Finally, Tripso’s capacity for data-driven GP discovery revealed a previously uncharacterized tissue-resident memory T cell GP with increased activity in atopic dermatitis. Its spatial co-localization with sebaceous gland-associated immune niches was demonstrated in spatial transcriptomic and proteomic data. Thus, by moving beyond single embeddings of cellular states, Tripso enables interpretable and actionable discoveries, demonstrating how GP-centric modelling can generate hypotheses with substantial biomedical relevance. By anchoring cellular representations in meaningful GPs, Tripso establishes a principled and biologically grounded framework towards the development of interpretable virtual cell models.

## Introduction

Cell identity, state and function are shaped by the coordinated expression of genes that reflect underlying biological processes during diverse contexts, such as differentiation, homeostasis and disease. Single-cell molecular profiling technologies enable us to capture up to tens of thousands of genes per cell. However, conventional analysis workflows rely on selecting highly variable genes and reducing dimensionality through linear methods such as PCA (*1*) or deep learning models including variational autoencoders (VAEs) (*2*, *3*). The resulting cellular transcriptome embeddings serve as the basis for clustering, annotation, and downstream biological interpretation. More recently, the increase in number and scale of single-cell datasets (*4*, *5*) together with the success of transformers and generative artificial intelligence in natural language processing (*6*, *7*) has enabled the emergence of single cell foundation models (*8–15*). While these models can capture relationships between the expression of multiple genes, they typically represent each cell as a single embedding that entangles multiple sources of biological variation. This hinders the study of complex experimental designs spanning multiple conditions such as *in vivo* and *in vitro* systems or disease states (*3*, *16*, *17*). Moreover, as efforts shift toward building comprehensive “virtual cells”, single-cell foundation models face a dual challenge: balancing model capacity with interpretability, and systematically translating predictive performance into mechanistically informative and experimentally testable biological hypotheses (*18*). Thus, despite their increased scale, single cell foundation models do not consistently translate into improved performance or interpretability relative to simpler approaches (*19–22*).

A complementary strategy is to represent each cell through multiple gene programs (GPs), defined as coordinated sets of genes whose expression reflects specific biological processes, such as signaling responses, transcription factor activity, or disease-associated states. By decomposing the transcriptome into interpretable units, GP-based modeling captures distinct axes of biological variation, enabling systematic comparison of shared and context-specific programs across tissues, conditions, and diseases. Importantly, this avoids overreliance on a single embedding and allows program-level differences to be resolved to individual gene contributions.

Existing GP-based approaches typically summarize each program as a scalar score or constrain it to a single latent dimension. For example, matrix factorization methods such as Spectra (*23*) and interpretable VAEs including VEGA (*24*) and Expimap (*25*) improve interpretability but limit programs to linear or one-dimensional representations. Other methods, such as DeepGSEA (*26*) or disentanglement frameworks (*2*, *27–29*), learn GP- or condition-specific representations but depend on supervision or predefined covariates, restricting their ability to capture context-dependent variation across highly divergent conditions, such as *in vivo* and *in vitro* systems.

To address these limitations, we developed Tripso (Transformers for learning Representations of Interpretable gene Programs in Single-cell transcriptOmics), a self-supervised framework that represents cellular state through multiple GP-specific embeddings. Tripso learns contextualized gene, GP and cell embeddings, quantifies gene-and GP-level contributions to GPs and cell identity respectively, and enables discovery of novel GPs. By combining the interpretability of GPs with the flexibility of transformer architectures, Tripso supports principled comparison of cellular states across tissues and conditions. In human hematopoiesis, Tripso resolved age-dependent shifts in GP activity. By integrating *in vivo* and *in vitro* hematopoiesis data, Tripso identified systematic differences between stem and differentiated cell states, and highlighted candidates to improve *in vitro* stem cell maintenance, which we validated experimentally. In a cross-disease dataset of human skin, Tripso uncovered disease-associated GPs that were independently validated using matched spatial transcriptomic and proteomic measurements. Overall, Tripso provides a scalable foundation for interpretable single-cell modeling that enables hypothesis generation and experimental optimization across development, disease, and engineered in vitro systems, and supports the development of interpretable virtual cell models.

## Results

### Self-supervised learning of gene program representations

Tripso models cell states as a hierarchy of interpretable representations: the first two compose the base model (gene and GP representations) and the final representation is for the global model (cell representation). First, a gene encoder generates contextualized gene embeddings *via* self-attention among genes within individual cells (*Gene representation*) (**Fig. 1a**). These embeddings capture context-dependent expression relationships between genes within a cell.

**Fig. 1:**
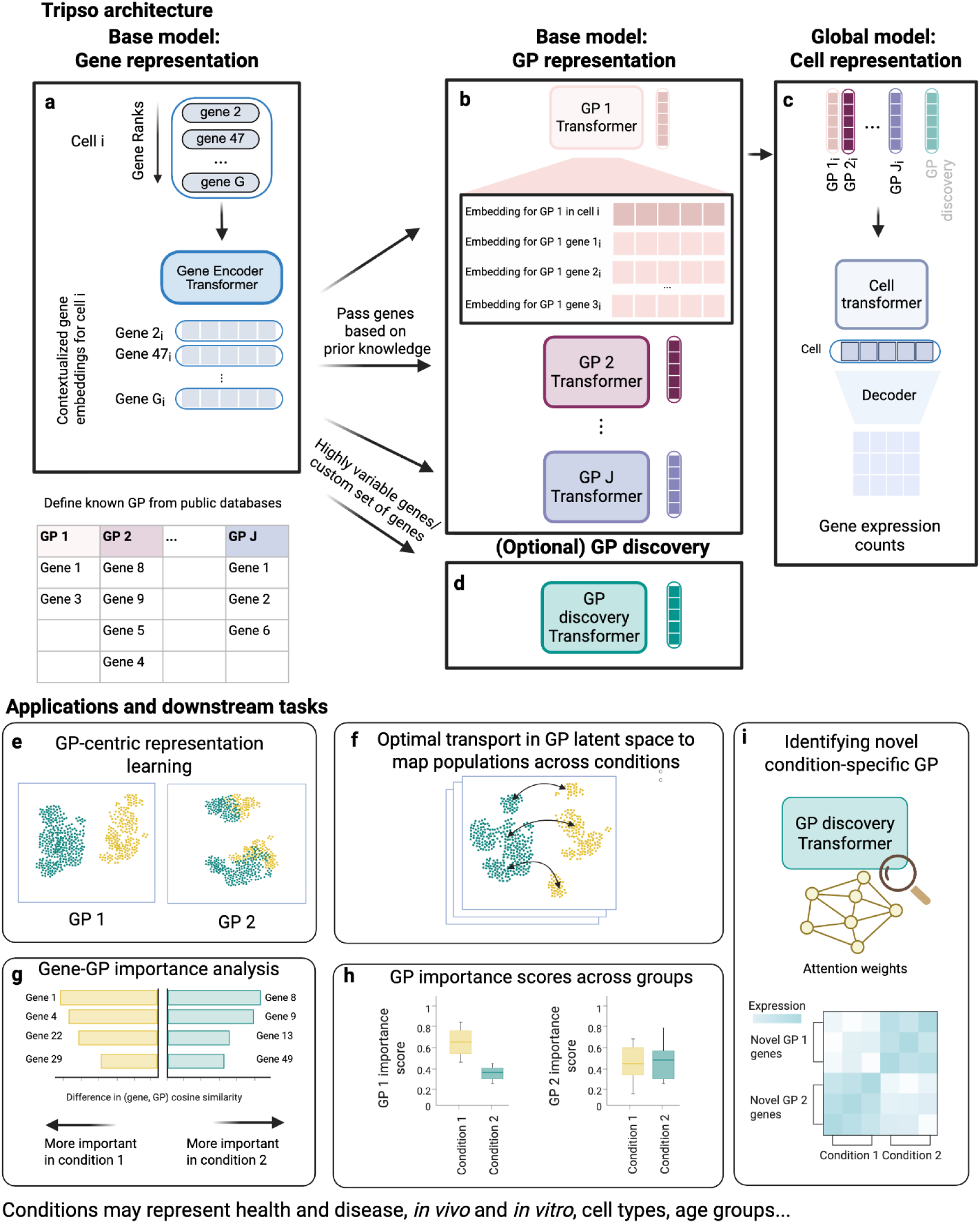
Self-supervised learning of gene programs using Tripso. (**a**) Tokenized scRNA-seq count data are used as input to a transformer model, which produces contextualized gene embeddings at single-cell resolution. Curated GPs are initialized from public databases. (**b**) GP-specific transformers learn latent representations for each GP based on their constituent genes. (**c**) A global cell block integrates GP representations to learn a unified cell-level embedding. (**d**) Optionally, an additional GP discovery transformer is defined using highly variable genes or a user-specified gene set to infer data-driven GPs. (**e**) UMAP visualization of GP CLS embeddings. (**f**) Optimal transport analysis performed in the GP latent space to infer mappings of cell populations across conditions. (**g**) Gene-to-GP cosine similarity is used to quantify the contribution and importance of individual genes to learned GP representations. (**h**) GP importance to the overall cell representation is quantified by systematic ablation of individual GPs and quantifying the resulting deviation in cell embeddings. (**i**) Clustering of gene-gene attention patterns from the GP discovery module defines novel, data-derived GPs, which can be interpreted by examining their expression patterns in the original data space.

The second component is a set of predefined GP transformer blocks. The relevant GPs are user-defined based on the biological question of interest, and the member genes are extracted from public databases (**Methods**). Although there is no strict upper limit on program size, very large gene sets reduce interpretability, and effective learning requires programs of sufficient size (**Methods**). The gene embeddings are routed to GP-specific transformers based on this predefined membership. Each GP block incorporates a special token (CLS) that attends to associated gene embeddings, acting as a nonlinear summary of the GP genes in each cell (*GP representation*) (**Fig. 1b**), and the GP blocks are trained using a masked-language modeling objective (**Methods**). Transformers provide a flexible architecture for modeling GPs, as they allow context-dependent weighting of genes within a program and accommodate both up- and downregulated expression patterns across diverse cellular states.

Finally, in the global model, a cell representation is learned by attending to the GP embeddings (*Cell representation*) (**Fig. 1c**). The GP embeddings are aggregated and passed to a cell decoder, which applies self-attention across GP embeddings to yield cell representations. This decoder is trained using a negative binomial loss to reconstruct original gene expression counts (**Methods**).

In addition to predefined GPs, Tripso can discover GPs directly from data, independent of existing annotations (**Fig. 1d,i**). In this setting, the transformer operates on a selected set of genes (for example, highly variable genes or a user-defined gene set), and the learned attention patterns capture gene-gene relationships within individual cells. These attention patterns can be interpreted in two complementary ways. First, genes can be ranked by their attention weights to identify those most relevant to specific cellular states. Second, genes with similar attention profiles can be grouped to define context-specific programs. This strategy leverages the premise that genes participating in shared GPs exhibit similar interaction patterns, enabling data-driven identification of GPs.

The GP-centric representation achievable by Tripso enables a wide range of downstream analyses, which are readily implementable using standard single-cell genomics tools. GP CLS embeddings can be visualized using UMAPs (**Fig. 1e**), or used to perform optimal transport analyses across conditions, such as between *in vivo* and *in vitro* data, while anchoring comparisons in a GP-specific latent space (**Fig. 1f**). To quantify the contribution of individual genes to a given GP representation, we compute the cosine similarity between each gene embedding and the corresponding GP CLS token. This provides a gene-level importance score within each program, enabling identification of which constituent genes drive GP activity in a given cell state and how their relative contributions shift across conditions (**Fig. 1g**). To assess the relative importance of individual GPs to the overall cell representation, we systematically ablate each GP by setting its embedding to zero and measuring the resulting change in the cell representation using cosine similarity. The magnitude of this deviation from the original representation defines a GP importance score (**Fig. 1h**). Data-derived GPs can be interpreted by examining their expression patterns in the original data or across complementary modalities, thereby revealing the cell states and biological contexts in which they are active (**Fig. 1i**).

### Benchmarking Tripso for gene program activity modeling

We benchmarked Tripso against representative methods from two widely used classes of approaches for modeling GP activity: matrix factorization-based models and deep generative models. Specifically, we compared Tripso to Spectra (*23*), which uses non-negative matrix factorization, and to Expimap (*25*), an interpretable variational autoencoder-based framework. We did not include another tool, DeepGSEA (*26*), in this benchmark, as it relies on supervised learning, whereas Tripso is designed for label-free, self-supervised analysis.

To compare each GP model’s ability to capture ground-truth GP activity, we used a large-scale Perturb-seq dataset (*30*), in which key regulators were genetically perturbed in six cancer cell lines stimulated by cytokines including TGFβ and TNFα (**Fig. 2, Methods**). For benchmarking, we focused on TNFα and TGFβ stimulation (**Methods**), for which curated transcriptomic response signatures are available in PROGENy (*31*). These signatures were used as input GPs for all methods, providing a well-defined reference for evaluating each method’s ability to discriminate stimulation-specific GP activity within the corresponding GP embedding space. After filtering, this dataset comprised 623,000 cells and 98 genetic perturbations. We designed two benchmark tasks to assess the quality of the learned GP representations. First, we tested whether GP embeddings could accurately distinguish cells based on their cytokine stimulation (i.e., distinguishing TGFβ-stimulated cells from those stimulated by other cytokines, based on TGFβ representation alone). Tripso outperformed existing methods, achieving an average increase of 0.15 in F1-score against the second best performing method (**Fig. 2c, Supplementary fig. S1a**).

**Fig. 2:**
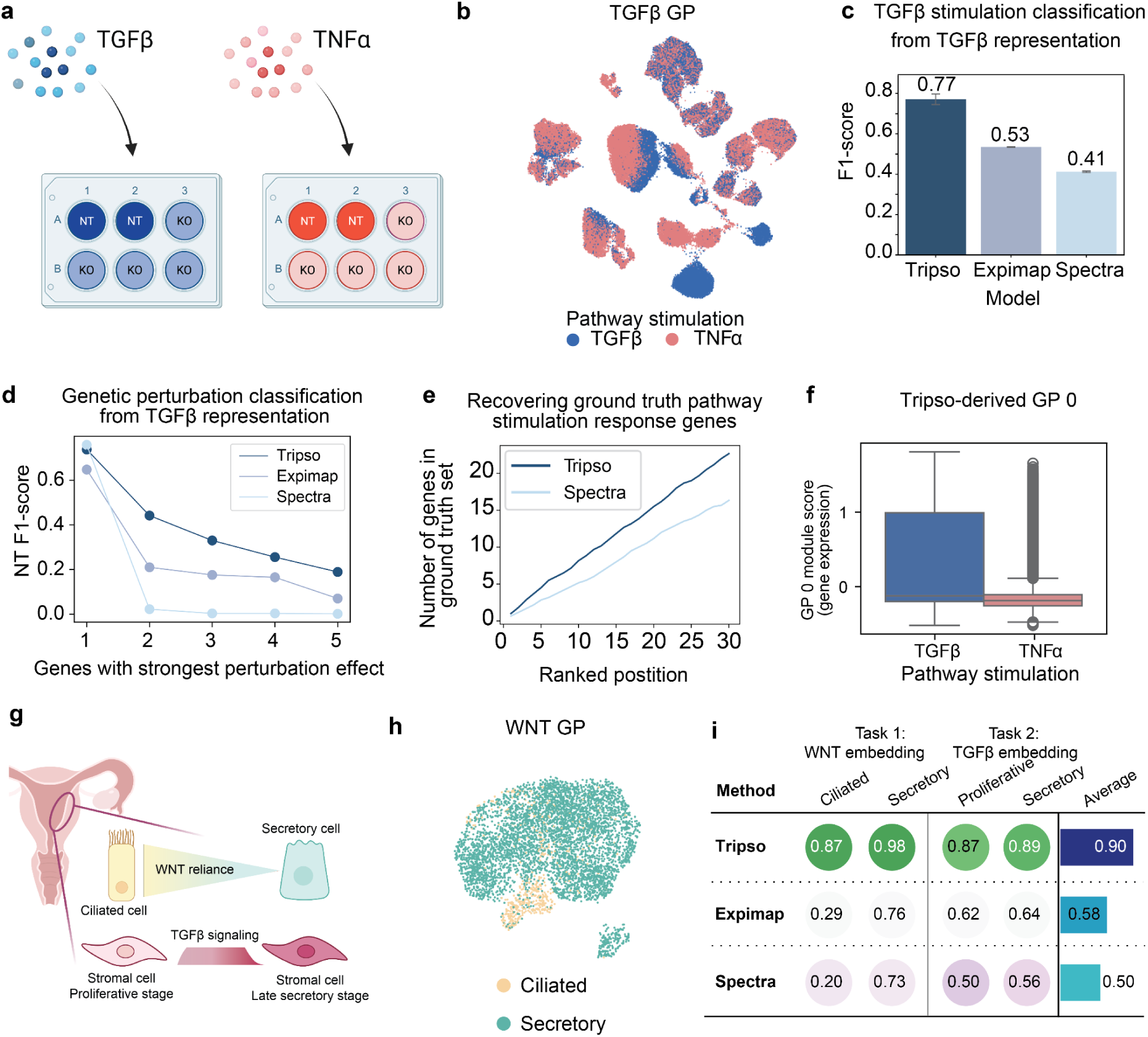
Benchmarking Tripso. **a**) Schematic overview of the filtered Perturb-seq dataset (*30*). NT - nontargeting control, KO - regulator knockdown. **b**) UMAP representation of TGFβ GP latent space. Each dot represents a cell, colored by the ground truth cytokine stimulation. **c**) F1-score for TGFβ classification from TGFβ GP representation learned by each model. Error bars represent variation across three random seeds. **d**) F1-score for classifying non-targeting controls from TGFβ GP representation learned by each model. The x-axis represents the number of perturbed gene classes. **e**) Number of genes previously reported as being involved in cytokine stimulation response in the original publication (*30*) recovered by Tripso and Spectra. The x-axis represents the ranked gene position as output by both methods, the y-axis represents the number of ground truth genes identified at each position. Random chance reflects the proportion of ground truth genes among the total available genes. **f**) Module score (computed on original gene expression space) for Tripso GP 0 in TGFβ- or TNFα-stimulated cells. **g**) Schematic representation of the endometrium and two examples of known patterns of GP activity (*32*). **h**) UMAP representation of learned WNT GP latent space. Each dot represents a cell, colored by broad cell type. **i**) Benchmarking results on the endometrium dataset. Each number corresponds to the average F1-score for the class of that column, based on the representation learned for the GP indicated above the relevant column. Color intensity represents the value (purple = lower, green = higher, blue intensity in the “Average” column represents ranking, darker = higher).

To further contextualize performance, we compared Tripso against two non-machine learning baselines. The first, Scanpy score genes, quantifies GP activity by averaging the expression of GP-associated genes in each cell relative to a reference gene set. The second baseline simply concatenates the log-normalized expression values of GP genes to define the GP embedding. In both cases, Tripso achieved superior F1-scores for identifying pathway stimulations (**Supplementary fig. S1b,c**). Importantly, representations derived directly from log-normalized expression are particularly sensitive to batch effects. Using scIB metrics (*16*), we confirmed that Tripso embeddings improved batch mixing relative to log-normalized gene expression alone (**Supplementary fig. S1d**).

We next assessed each method’s ability to detect genetic perturbations, which typically induce weaker responses than cytokine stimulations (*30*). Specifically, we evaluated how well each method could discriminate cells perturbed by a genetic knock-out from non-targeting controls, considering cells stimulated with the relevant pathway cytokine for each task. To quantify the difficulty of the task, we used the average gene-level perturbation effect score reported in the original publication (*30*). Tripso consistently outperformed other methods in classifying genes with the strongest perturbation effects (**Fig. 2d, Supplementary fig. S1e,** standard deviations in **Supplementary table t1**). Across all methods, performance decreased as the classification task expanded to include perturbations with progressively weaker transcriptomic effects, reflecting the subtler and more variable phenotypes induced by these perturbations. We also evaluated the ability of the learned cell representation from each method to discriminate cytokine perturbation status. Tripso maintained strong discriminative performance at the cell level, outperforming alternative approaches (**Supplementary fig. S1f**).

Finally, we leveraged the pathway-response genes defined in the original publication (*30*) to evaluate each method’s ability to recover class-specific GPs, corresponding to the discovery of novel pathway-specific signatures. In this benchmark, prior knowledge of one pathway signature (TGFβ or TNFα) was provided while the target GP was withheld, and performance was assessed on recovery of the held-out signature. Averaged over three random seeds and across both tasks, Tripso prioritized more stimulation-response genes within the top 15, with an increase of 3.8 hits on average compared to Spectra (48% relative improvement; **Fig. 2e**; per-task means and standard deviations in **Supplementary Table t2**). In this setting, Spectra learned one (and occasionally two) latent factor per class, typically resulting in GPs comprising hundreds of genes with non-zero weights, complicating biological interpretability. In contrast, by clustering correlations of gene attention scores, Tripso delineated biologically meaningful structured GPs that captured both cell-type and pathway stimulation effects (**Fig. 2f; Supplementary fig. S1g,h**). Robustness was assessed by repeating the full GP discovery pipeline across multiple random initializations, including both model training and clustering (**Supplementary fig. S1i**). This illustrates that Tripso’s GP discovery setting can uncover meaningful and reproducible GPs.

Having established Tripso’s performance in controlled perturbation experiments, we next evaluated its ability to resolve biologically meaningful GP activity in a heterogeneous, real-world tissue setting. We applied Tripso to the human endometrial cell atlas (*32*), which contains transcriptomic data from 313,527 cells, spanning 36 cell types (**Fig. 2g**). Input GPs were defined using pathway signatures from PROGENy (*31*) (**Methods**). We focused on two biological scenarios with known differences in GP activity. First, epithelial cell differentiation into ciliated or secretory cells is controlled by the balance of WNT and Notch signalling, and WNT inhibition leads to depletion of ciliated cells (*33*). Tripso’s WNT embeddings (Task 1) (**Fig. 2h**) improved discrimination between these two cell states compared to Expimap and Spectra, achieving a mean per-class improvement of 0.40 in F1-score over the second best performing method (computed as the average of the per-class F1-score differences; **Fig. 2i**, standard deviations across 3 random seeds reported in **Supplementary table t3**). Second, we examined TGFβ signalling in mesenchymal cells (Task 2), which shows phase-specific regulation across the menstrual cycle, including decreased TGFβ ligand expression and increased inhibitor expression in the late secretory phase (*32*). Tripso embeddings achieved an average F1-score of 0.88 in discriminating menstrual cycle stages, compared to 0.63 and 0.53 for Expimap and Spectra, respectively (**Fig. 2i**, standard deviations across 3 random seeds reported in **Supplementary table t3**).

Taken together, these benchmarks indicate that Tripso’s embedding-based representation of GP activity provides greater discriminative capacity than a single scalar value.

### Tripso identifies age-specific patterns in human hematopoiesis

We first investigated whether Tripso could resolve age-specific GP activities underlying human hematopoiesis, a highly dynamic system that changes across the human lifespan, particularly during development and in ageing. Beginning prenatally in sequential and overlapping waves from the yolk sac and fetal liver, hematopoiesis within bone marrow is established by the end of the first trimester, with bone marrow becoming the primary site of hematopoiesis postnatally (*34*). Age-specific lineage differentiation biases are observed prenatally and further amplified during human ageing (*35–39*). These phenomena are underpinned by time and lineage-dependent GPs, providing an ideal setting to test Tripso’s exploratory capabilities.

We compiled a corpus of human hematopoietic scRNA-seq datasets (**Fig. 3a** and **Methods**). First, we newly generated a CITE-seq dataset of 79,583 blood and immune mononuclear cells (MNC) with paired transcriptome and surface protein measurements from age-specific hematopoietic tissues (yolk sac, fetal liver, and bone marrow) across the human lifespan (ranging from 6 post conception weeks to 91 years of age) (**Methods, Fig. 3b and Supplementary fig. S2a-c,** hereafter MNC dataset). We further combined this with a recently generated human CD34^+^ hematopoietic reference atlas (*40*) (**Fig. 3c**, hereafter CD34^+^ dataset). Together, these datasets encompass the full spectrum of human hematopoietic cells spanning from the most immature hematopoietic stem cells (HSCs) to mature populations across eight lineages: megakaryocyte, erythroid, eosinophil/basophil/ mast cell, B-lymphoid, natural killer/T-lymphoid, monocyte/macrophage, dendritic cell and plasmacytoid dendritic cell (**Supplementary table t4**). We further augmented our training set with seven additional publicly available scRNA-seq datasets profiling healthy hematopoiesis across stages and tissues (*39*, *41–47*). Additionally, we included two distinct *in vitro* hematopoiesis datasets derived from human cord blood cultured under differentiation (*48*) and maintenance conditions (*49*) to compare with our human *in vivo* data. Collectively, our hematopoietic single-cell data corpus included 498,836 cells from 98 healthy donors and 46,667 cells from in *vitro* culture experiments. To train Tripso, we selected a set of well-established hematopoietic transcription factors and their target genes from CollecTRI (*50*) and complemented this list with general cytokine response gene sets from PROGENy (*31*) (**Methods,** GP curation). In this section, all GPs named after a transcription factor consist of the target genes attributed to that transcription factor in CollecTRI (*50*)

**Fig. 3:**
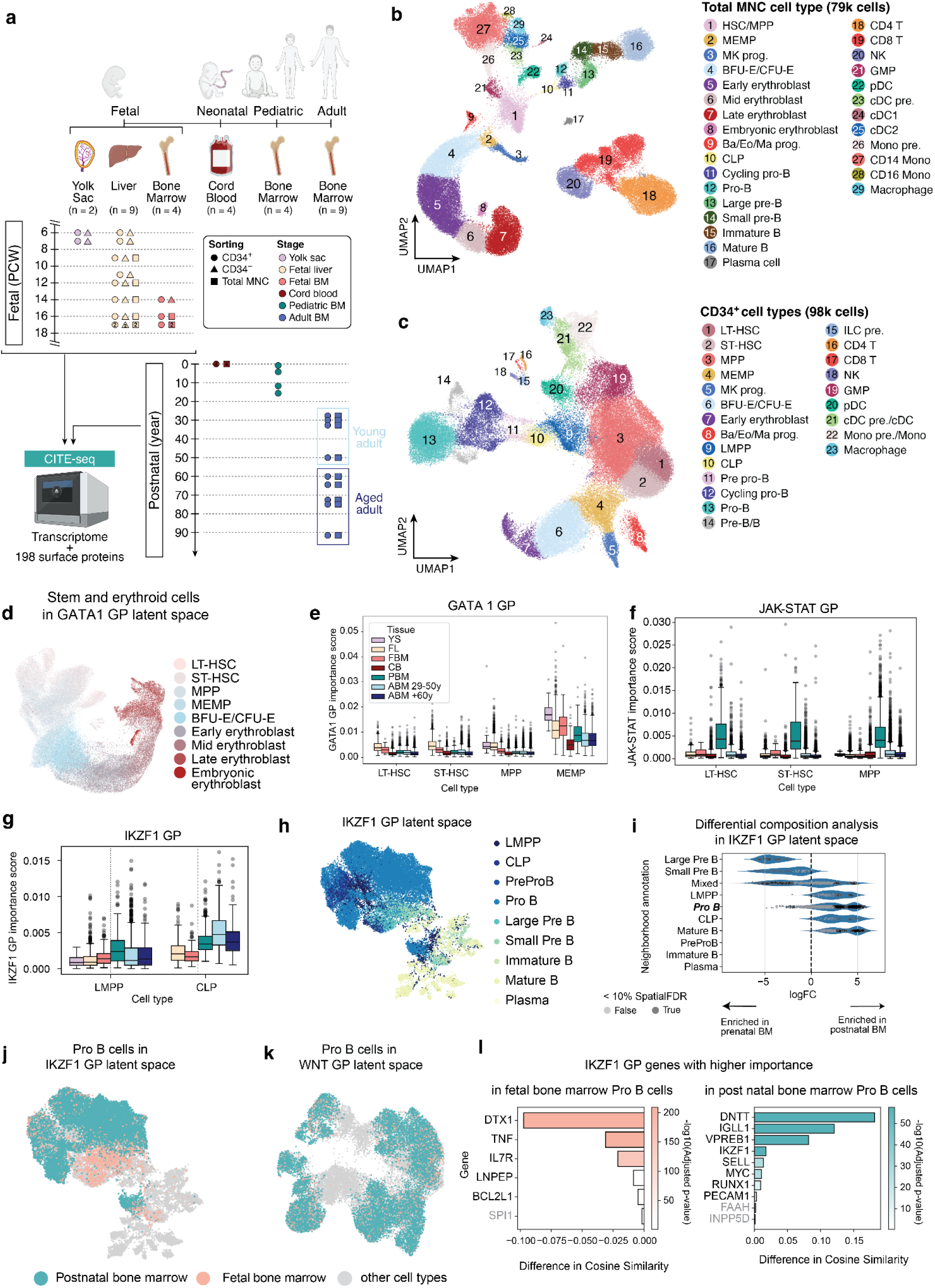
Tripso identifies age-specific cell states in hematopoiesis. **a**) Schematic describing tissue of origin and patient age for in vivo datasets of interest. (**b/c**) UMAP of the unsorted MNC (**b**) and CD34+ enriched (**c**) datasets. Cells are colored by cell type, UMAP and cell type annotation are based on transcriptomic and surface proteomic data. **d**) UMAP of the Tripso GATA1 GP embedding space, including stem, progenitor and erythroid cells from the CD34+ and MNC datasets, colored by cell type. **e**) GATA1 GP importance scores in stem and progenitor cell populations, separated by tissue. **f**) JAK-STAT GP importance scores in HSPC populations, separated by tissue (colors as in panel e). **g**) IKZF1 GP importance scores in LMPPs and CLPs, separated by tissue (colors as in panel e). Tissue categories with less than 10 cells per value are removed from visualization in panels e, f and g. GP importance scores in e, f and g represent averages across 3 model runs initialized with 3 random seeds. **h**) UMAP of bone marrow (prenatal, pediatric, adult) B lineage cells in IKZF1 GP embedding space, colored by cell type. **i**) Milo differential composition analysis results, testing for neighborhoods enriched in postnatal bone marrow (PBM, ABM) vs fetal bone marrow. Darker circles represent spatial FDR < 10%. In (i) and (h), Pro-B and Cycling Pro-B cells are grouped together. **j**) UMAP of bone marrow B lineage cells in IKZF1 GP embedding space, pro-B cells are highlighted and colored by tissue stage. **k**) UMAP of bone marrow B lineage cells in WNT GP embedding space, pro-B cells are highlighted and colored by tissue stage. **l**) Differential gene importance for IKZF1 GP in pro-B cells in postnatal (ABM, PBM) compared to fetal bone marrow. Gene names in gray correspond to differences with adjusted p-values >0.05. Gene cosine similarity scores represent averages across 3 model runs initialized with 3 random seeds. YS - yolk sac, FL - fetal liver, FBM - fetal bone marrow, CB - cord blood, PBM - pediatric bone marrow, ABM - adult bone marrow, LT-HSC - long term HSC, ST-HSC - short term HSC, MPP - multipotent progenitors, MEMP - megakaryocyte-erythroid-mast cell progenitors, LMPP - lymphoid multipotent progenitors, CLP - common lymphoid progenitors.

We first validated that Tripso representations captured biologically meaningful patterns in hematopoiesis, focusing on well-established patterns in each lineage. In the erythroid lineage, we focused on the GATA1 GP which contains GATA1 target genes, as it is a canonical erythroid transcription factor, and well-established regulator of red blood cell differentiation (*51*). We projected HSCs, multipotent progenitors (MPPs) and erythroid cells from our in-house CD34^+^ and MNC datasets into the GATA1 GP embedding space (**Fig. 3d**). The resulting embedding recapitulated expected erythroid lineage cell states, with cells clustering according to lineage progression from HSCs through intermediate progenitors to late erythroblasts. We then turned to the myeloid and megakaryocyte lineages, focusing on RUNX1, which is crucial for both monocyte (*52*) and megakaryocyte (*53*) differentiation. Consistent with this role, visualization of the first two principal components of the RUNX1 GP latent space shows that monocytes and megakaryocytes clustered separately from precursor, progenitor and erythroid cells along two differentiation trajectories (**Supplementary fig. S2d**). Projection of the monocyte surface marker CD14 from CITE-seq data followed this pattern (**Supplementary fig. S2e**), and RUNX1 importance scores were highest in granulo-monocyte progenitors (GMPs) and megakaryocytes **(Supplementary fig. S2f**). GP importance scores also matched expected cell type-specific patterns. For example, *FLI1* is required for megakaryocyte differentiation (*54*), and FLI1 GP importance score was highest in megakaryocytes (**Supplementary fig. S2g**). Together, these analyses demonstrate that the main axes of variation in GP embeddings capture expected and biologically meaningful patterns of hematopoiesis.

Having validated Tripso’s representations against well-established biological patterns, we next applied the model to age-specific variation in hematopoiesis. To do this, we explored age-dependent patterns observed in the immature HSC/multipotent progenitor (MPP) fraction. GATA1 GP importance was elevated in fetal compared to adult HSC/MPPs and reached highest levels in fetal megakaryocyte-erythroid-mast cell progenitors (MEMPs), consistent with the increased erythroid output during fetal hematopoiesis (*35*) (**Fig. 3e**). Furthermore, increased JUNB GP importance was observed in older age groups (adult bone marrow samples) as previously reported (*55*, *56*) (**Supplementary fig. S2h**).

In addition to these expected age-dependent GP importance, pediatric HSC/MPPs (ages 1 to 16) exhibited significantly higher importance of the JAK-STAT GP (**Fig. 3f, Supplementary fig. S3a**), and this effect was consistent across pediatric donors (**Supplementary fig. S3b**). Since the JAK-STAT pathway is involved in multiple cytokine and growth factor signals (*57*), we next explored the gene-level importance within JAK-STAT GP. Genes with elevated importance within pediatric HSC/MPPs significantly overlapped with type I interferon response genes (gene set enrichment analysis adjusted p-value < 10^-10^, **Supplementary fig. S3c,d**). This pediatric specific interferon-JAK-STAT activity is evocative of the transient interferon pulse promoting perinatal HSPC expansion observed in mice (*58*), suggesting a potentially conserved mechanism underlying postnatal HSPC expansion in humans (*59–61*).

Next, to investigate age-associated differences in hematopoietic lineage differentiation, we focused on the B cell lineage, where differences between fetal and adult lymphopoiesis have been previously reported (*62*, *63*). We first examined GPs displaying significant differences in importance scores between prenatal and postnatal tissue lymphoid progenitor populations. Cord blood was excluded from the primary prenatal-postnatal comparison because it represents a perinatal circulating compartment. Within this framework, IKZF1, a critical transcription factor for lymphoid lineage specification (*64*), emerged among the top GPs enriched postnatally in lymphoid multipotent progenitors (LMPP) and common lymphoid progenitors (CLP) (**Fig. 3g**, **Supplementary fig. S3e**).

To investigate the implications of this shift for B-lineage differentiation, we explored the IKZF1 GP embedding space. Differential abundance testing using Milo (*65*, *66*), comparing postnatal bone marrow (ABM and PBM) to fetal bone marrow (FBM), revealed distinct Pro-B cell neighborhoods enriched either prenatally or postnatally within the IKZF1 GP latent space (grouping Pro-B and cycling Pro-B cells together, **Fig. 3i,j**). Importantly, this effect was specific to the IKZF1 GP embedding. As a control, we performed the same Milo-based differential abundance analysis using the WNT GP, which showed no significant prenatal-postnatal difference in GP importance scores and the smallest fold-change in IKZF1 GP importance between pre– and postnatal lymphoid progenitors of our included GPs (**Supplementary fig. S3f**). In the WNT latent space, prenatal and postnatal pro-B cells largely overlapped (**Fig. 3k**), and no differential enrichment was detected at FDR<10%. Thus, the observed divergence of pro-B cell states is not a general feature across all GP embeddings but reflects a program-specific developmental effect captured by the IKZF1 GP.

To understand the genes driving these differences, we examined IKZF1 GP genes that were differentially important between prenatal and postnatal pro-B cells. Analyses were performed both on all pro-B cells combined (**Fig. 3l**) and separately for cycling and non-cycling subsets, yielding consistent results (**Supplementary fig. S3g,h**). In line with the reciprocal control of IL7R versus pre-BCR-driven programs that prevents concurrent proliferation and recombination to limit genomic instability (*67*), prenatal pro-B cells favored expansion, expressing higher *DTX1* (promoting B-lineage commitment over T-cell differentiation (*68*)), and increased *BCL2L1* (an anti-apoptotic IL7R target (*69*)(*70*)). By contrast, postnatal pro-B cells emphasized BCR diversification, with increased importance of *IGLL1/VPREB1* (pre-BCR components) and *DNTT* (mediating N-nucleotide addition for increasing BCR diversity (*71*) and previously reported to be upregulated in adult B-lymphopoiesis compared to fetal (*63*)). Repeating the analysis by directly comparing postnatal bone marrow pro-B cells with fetal liver pro-B cells produced consistent results, both in terms of differential abundance (**Supplementary fig. S3i,j**) and gene-level importance (**Supplementary fig. S3k,l**). Together, these results demonstrate that developmental differences in early B-lineage differentiation are captured in a program-specific manner within the IKZF1 GP embedding space and reflect a broad shift from prenatal proliferation to postnatal diversification.

Collectively, these findings demonstrate several key advantages of GP embeddings. First, Tripso embeddings enable the application of single-cell analytical methods designed for embeddings rather than scalar values, such as differential composition analysis. Second, they provide refined resolution of cell states that goes beyond standard cell-type classifications. Lastly, Tripso facilitates context-specific refinement and interpretation of GP representations.

### Tripso-guided mapping and prioritization of perturbations for *in vitro* hematopoietic cells

Accurate *in vitro* models of human cells are essential for biological discovery, drug screening, and cell-based therapies. The importance of *in vitro* human models is further underscored by the fundamental differences between mouse and human hematopoiesis (*72*). Although recent advances have enabled longer-term HSC maintenance in culture (*73*), the extent to which these cells preserve *in vivo* stem cell molecular programs remains unclear. Optimization of *in vitro* protocols is further constrained by the vast combinatorial space of perturbations and the absence of a quantitative, high-dimensional reference for evaluating cell-state improvements. Thus, we sought to leverage Tripso as a means to systematically map *in vitro* HSCs to their *in vivo* counterparts and prioritize interventions that improve *in vivo* relevance.

To validate the GP-resolved framework in an *in vitro* setting, we analyzed a published dataset in which cord blood-derived CD34^+^ HSPCs were differentiated *in vitro* over 17 days (*48*) (**Fig. 4a**). As expected under these culture conditions, erythroid differentiation was dominant, and we therefore focused on this lineage. Comparing GP importance scores across the *in vivo* erythroid trajectory reconstructed from our CD34⁺ and MNC datasets described above to those derived from the *in vitro* data revealed broadly concordant dynamics: VEGF GP importance was highest early, consistent with its role in HSC survival (*74*) and early erythroid differentiation (*75*, *76*), whereas the LMO2 GP, associated with later erythroid differentiation (*77*), gained importance terminally (**Fig. 4b,c**).

**Fig. 4:**
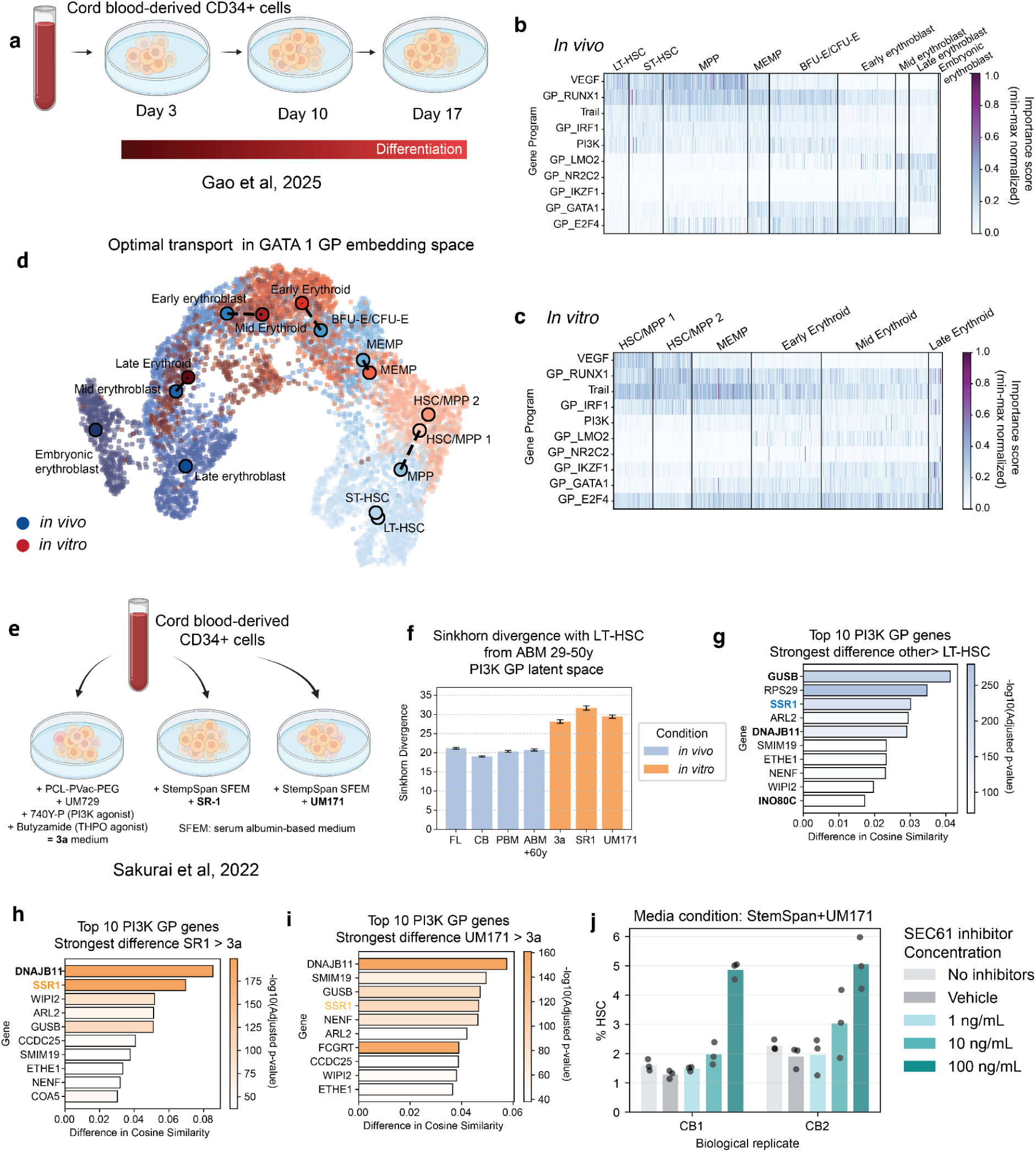
Gene program-guided *in vivo–in vitro* mapping and perturbation prioritization in hematopoiesis. **a)** Overview of experimental data generation for previously published cord blood in vitro differentiation dataset (*48*). (**b/c**) Heatmap showing min-max normalized GP importance score by erythroid cell type *in vivo* **b**) and *in vitro* **c**). Each row represents a GP, the top/bottom 5 correspond to GPs whose importance scores decrease/increase the most as a function of differentiation. Each value reflects the importance score for an individual cell, grouped by cell types. Importance scores are obtained by averaging results across 3 Tripso model training runs initialized with 3 random seeds. **d**) UMAP of GATA1 GP latent space for *in vivo* and *in vitro* erythroid cells. Small dots represent individual cells, colored by cell type. Large labelled dots represent the real cell nearest to the cluster centroid for that cell type. Dashed lines represent pairings computed by unbalanced optimal transport. **e**) Overview of experimental data generation for previously published stem cell maintenance dataset (*49*). **f**) Barplot showing the Sinkhorn divergence between cell populations described on the x-axis and LT-HSCs from adult bone marrow of patients 29-50 years old. Error bars indicate the standard deviation across 50 bootstrap iterations (750 cells sampled per iteration). **g**) Differential gene importance for PI3K GP genes in stem and progenitor cells compared to LT-HSC compared *in vivo.* **h**) Differential gene importance for PI3K GP genes in HSPCs cultured in SR-1 media compared to 3a media. **i**) Differential gene importance for PI3K GP genes in HSPCs cultured in UM171 media compared to 3a media. Gene cosine similarity scores represent averages across 3 model runs initialized with 3 random seeds. Gene names in bold correspond to genes which were significantly differently expressed (adjusted p-value <0.05) in pseudobulked differential gene expression. **j**) Proportion of immunophenotypic HSCs (CD34+ CD45RA− CD90+ EPCR+, as measured by flow cytometry) detected under different concentrations of SEC61 inhibitor or control (no inhibitors and vehicle). Each cord blood (CB) represents a biological replicate, each dot represents a technical replicate.

We next examined GATA1, whose GP importance displayed context-specific behavior: it declined along the *in vivo* trajectory but remained elevated throughout *in vitro* differentiation. Some divergence between *in vivo* and *in vitro* trajectories was expected, as GlyA⁺ late erythroblasts had been excluded from the *in vitro* dataset (*48*). To assess this computationally, we applied unbalanced optimal transport in the GATA1 GP embedding space to map *in vitro* cells to the *in vivo* reference (**Methods**). Early and mid-erythroid populations aligned across conditions, while the terminal *in vivo* erythroblasts remained unmatched, recapitulating the truncated trajectory in the *in vitro* protocol, without prior knowledge of the experimental design (**Fig. 4d**). This confirms that the GP-resolved framework supports biologically faithful, imbalance-aware alignment of *in vivo* and *in vitro* trajectories and cell states.

Building on this ability to compare *in vivo* and *in vitro* cells, we next applied Tripso to identify candidate gene targets to improve *ex vivo* HSC maintenance. HSCs are of considerable therapeutic importance for stem cell transplantation and gene therapy, but their low abundance necessitates robust *in vitro* expansion strategies. To analyze molecular differences between culture conditions, we leveraged a published scRNA-seq dataset (*49*) (**Fig. 4e**), which established a cytokine-free medium termed ‘3a’ that improved maintenance of functional human HSCs *in vitro*, as demonstrated by increased phenotypic HSC numbers and enhanced transplantation capacity. The scRNA-seq dataset includes cells cultured with 3a media and two baseline media conditions, with StemSpan supplemented with small molecules SR-1 or UM171. We applied Tripso to these scRNA-seq profiles to compare GP activity across culture conditions and relative to *in vivo* reference HSC states.

We first identified GPs important for HSC identity *in vivo* using the CD34+ reference dataset introduced above. Among the most distinctive was PI3K (**Supplementary fig. S4a).** Notably, the 3a medium used to culture cells in the published scRNA-seq dataset (*49*) contains the PI3K agonist, 740 Y-P. This, in addition to previous work implicating PI3K in regulation of HSC maintenance (*78*), prompted us to focus our subsequent analyses on the PI3K GP.

To assess similarity between *in vitro* and *in vivo* HSCs, we projected cells into the PI3K GP embedding space, thus ensuring that comparisons specifically reflected alignment within this biologically relevant program. We quantified differences between cell populations using Sinkhorn divergence, a distribution-level metric that captures structural differences between populations (**Methods**). To contextualize these Sinkhorn divergence values, we reported divergences among the most immature phenotypic long-term HSCs (LT-HSCs) from different tissues relative to adult bone marrow LT-HSCs (29–50 years). In the PI3K GP embedding space, the *in vitro* cells fell outside of the range observed across human tissues. Among the *in vitro* conditions, cells cultured in 3a medium showed the smallest divergence from adult bone marrow LT-HSCs (**Fig. 4f**), consistent with the higher reconstitution capacity of 3a-cultured HSCs (*49*).

To pinpoint genes within the PI3K GP whose modulation could enhance HSC maintenance in culture, we next performed a gene-level importance analysis. Across both *in vivo* and *in vitro* datasets (including SR-1 and UM171 media conditions), we found that the *SSR1* gene was consistently more important in less stem-like states (**Fig. 4g-i**). Its expression increased progressively along differentiation trajectories across all lineages (**Supplementary fig. S4b**), linking low *SSR1* expression to the stem-like state rather than any specific lineage. *SSR1* encodes a component of the translocon-associated protein (TRAP) complex, which interacts with the Sec translocon in the endoplasmic reticulum (ER) to support protein biogenesis (*79*). Interestingly, *SEC61G,* encoding for the Sec61-γ subunit and included within the hypoxia GP, displayed a similar pattern: higher gene-level importance in less stem-like states in both *in vivo* and *in vitro* datasets (**Supplementary fig. S4c–e**), converging on ER translocon machinery as a candidate regulator of hematopoietic differentiation.

To experimentally test Tripso’s prediction that SEC61 components are upregulated during differentiation, and that their inhibition may help maintain cells in a more stem-like state, we treated cord blood-derived CD34+ cells with the SEC61 inhibitor SEC61-IN-1 (TargetMol Chemicals Inc.) across the culture conditions described by Sakurai et al. Treatment with SEC61-IN-1 increased the frequency of immunophenotypic HSCs (CD34+ CD45RA− CD90+ EPCR+), specifically in media supplemented with UM171 and SR-1 (**Fig. 4j, Supplementary fig. S4f-h,** gating strategy provided in **Supplementary fig. S5**). In UM171 media, the median proportion of immunophenotypic HSC increased to 5.0% at 100 ng/mL, compared with 1.4% in the vehicle control. SEC61 inhibition was also associated with reduced production of CD14⁺ monocytes in UM171- and SR-1-treated cultures, and decreased CD235a⁺ erythroid cells across media conditions, as assessed by flow cytometry (**Supplementary fig. S6**). In contrast, no increase in immunophenotypic HSC proportion was observed in 3a medium (**Supplementary fig. S4g**). This outcome is consistent with our analysis design, which prioritized genes that were differentially important in UM171- and SR-1-containing conditions relative to 3a. Importantly, although pseudobulk differential expression analysis identified statistically significant differential expression of SSR1 (log fold change LT-HSC vs others: −0.25; adjusted p-value = 0.004) and SEC61G (log fold change LT-HSC vs others: −0.40; adjusted p-value < 0.001), these genes would have been difficult to prioritize using this conventional approach due to their small effect sizes. By contrast, Tripso’s GP-resolved approach nominated these SEC61 components as top candidates within the PI3K GP, directly guiding experimental validation. Together, these results identify SEC61 inhibition as a candidate strategy for increasing phenotypic HSC frequency *in vitro*, and illustrate how GP-based analyses can guide the refinement of stem cell culture protocols.

### Tripso discovers disease-associated gene programs in human skin inflammation

Finally, we sought to leverage Tripso to perform discovery of GPs across disease contexts. We therefore applied Tripso to human skin: the primary barrier protecting from environmental insults or infections. This barrier is disrupted in chronic cutaneous inflammatory disorders, such as atopic dermatitis (AD) and psoriasis, leading to substantial morbidity. Common to both AD and psoriasis is T helper-mediated inflammation, driven by dysregulated immune-epithelial crosstalk that promotes keratinocyte activation and tissue remodeling (reviewed in (*80*, *81*)). Despite partially overlapping T cell states in both diseases (*82*), they are characterised by distinct cytokine axes, with type 2 inflammation featuring IL-4 and IL-13 signalling in AD, and Th17 responses featuring IL-23 and IL-17 in psoriasis (*82*, *83*). Targeted therapies exploiting these cytokine axes, including those inhibiting type 2 cytokines JAK-STAT signalling or IL-17-mediated inflammation, exist, yet many patients with inflammatory skin disorders fail to achieve durable remission (*84*) or develop paradoxical skin reactions (*85*).

To identify shared and disease-specific transcriptional programs driving inflammatory skin pathology, we trained Tripso on a large single-cell atlas of human skin biopsies (*86*) (∼1.7 million cells across 338 samples; **Fig. 5a**) spanning healthy skin (830,885 cells) and 14 disease contexts, including atopic dermatitis (323,832 cells) and psoriasis (352,019 cells) as well as additional inflammatory, autoimmune and neoplastic condition (cell counts per condition **Supplementary table t5**). We then focused downstream analyses and validation on AD and psoriasis. We used matched spatial transcriptomics and proteomics to map disease-associated GPs to their tissue niches, localized tissue regions with distinct cellular composition and inflammatory features, as defined in the atlas (*86*). For model training, Tripso GP blocks were defined using transcriptional response signatures described in PROGENy (*31*). The Tripso GP discovery module (**Methods**) was restricted to Xenium Prime 5K Human Pan Tissue & Pathways Panel genes, enabling direct downstream validation in spatial transcriptomics data. GPs were inferred from patterns of gene-cell attention using two complementary strategies: lineage-conditioned discovery within specific cell types (i.e., performing the analysis on subsets of cells corresponding to the broad cell types defined in the atlas), and dataset-wide discovery across all cells, yielding multilineage GPs (MGP, **Methods**). We assigned each inferred GP a unified numeric identifier (GP1–GPN).

**Fig. 5:**
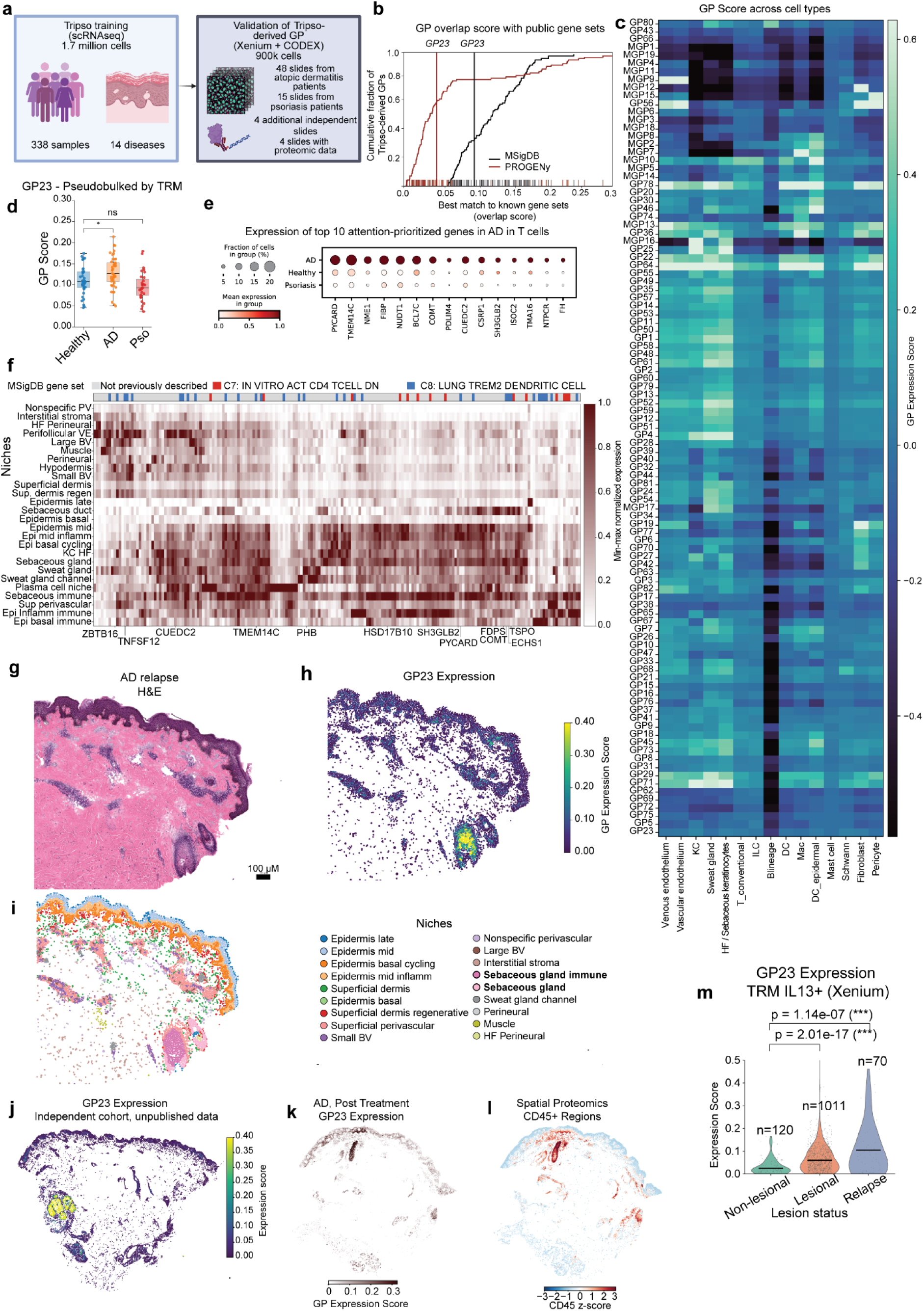
Tripso discovers new GPs associated with inflammatory skin conditions. **a**) Overview of computational set-up for GP Discovery in skin inflammatory diseases, encompassing training phase (left) and validation of GP (right). **b**) Empirical cumulative distribution of the maximum Jaccard overlap between each Tripso-derived GP and curated gene sets from PROGENy (GP used in Tripso base model, black line) or MSigDB datasets (union of C2, C7, C8 collection, red line). For each GP, we computed the maximum Jaccard index across gene sets in PROGENy or MSigDB and plotted the fraction of GPs with best-match overlap ≤ x. Tick marks indicate individual GP best-match scores; vertical lines highlight GP23. **c**) Heatmap showing the mean normalised expression of GPs and MGPs (rows) across cell types (columns) in scRNA-seq data. Hierarchical clustering is applied to rows and columns. **d**) Boxplot showing the aggregated expression of GP23 across healthy, AD and psoriasis (Pso) samples. Each dot corresponds to the mean normalised expression of GP23 across a single Xenium sample in TRM (annotated as either IL13+ or IL17+) cells only. Welch’s t test, p-val<<0.05. **e**) Dot plot of attention-prioritized genes in T cells across conditions (AD, healthy, psoriasis). Columns show the top 15 genes (selected by significance with logFC > 1) with increased attention in atopic dermatitis T-cells compared to other conditions; rows indicate condition. Each dot represents the mean expression in T cells within each condition, min-max scaled to 0-1 per gene across conditions and dot size indicates the fraction of T cells expressing the gene. **f**) Heatmap of GP23 gene-set expression across spatially annotated niches in the full Xenium cohort (60 sections). Columns indicate GP23 genes and rows indicate niche categories; both axes are hierarchically clustered. Top annotations mark genes that overlap the best-matching MSigDB C7 (immune perturbations) or C8 (cell-type signatures) gene sets. Bottom annotations highlight genes with established links to T cell biology. **g**) Haematoxylin-eosin (H&E) staining of the same AD lesional relapse section, as in panel g, highlighting a lymphocytic aggregate adjacent to the sebaceous gland niche. Scale bar, 100 μm. **h**) Spatial expression of GP23 in the same section (panels g-i). **i**) Same section as in panel g, colored by niche annotation. **j**) Spatial expression of GP23 module scores in an independent cohort, representative of in-house unpublished data for AD for this study. **k**) Spatial expression of GP23 module scores for a case of AD, Week 12 post Dupilumab initiation (same as panel j). Expression patterns are present for the epidermis and in the sebaceous gland. **l**) Spatial quantification of CD45 protein signal in the same section, computed from PTPRC (CD45) immunostaining pixel intensities and displayed as z-scored values. **m**) violin plots showing the expression of GP23 module scores in IL13+ TRM cells (Xenium), stratified by disease state in AD. “Non-lesional” denotes never-lesional skin; “Lesional” denotes baseline lesional skin; “Relapse” denotes week 8 after Dupilumab withdrawal. Welch’s t test between healthy and disease states, p-val < 0.05. Epi: Epidermis, Inflamm: Inflammation, Sup. Superficial, KC: keratinocyte, ILC: Innate lymphoid cell.

Comparison to curated resources, including PROGENy and MSigDB (*87*), indicated that most Tripso-derived GPs did not exhibit one-to-one correspondence with existing gene sets (**Fig. 5b, Supplementary fig. S7a**). Inferred GPs could not be fully reconstructed from existing gene sets (**Supplementary fig. S7b**), indicating that Tripso captured novel gene combinations spanning multiple annotation collections. Out of 101 inferred GPs, 6% were multilineage programs, whereas the remaining 94% were lineage-conditioned. Notably, GPs from both approaches showed coordinated expression patterns across multiple cell types, pointing to transcriptional processes that extend beyond canonical cell-type markers (**Fig. 5c**).

We then used Tripso-derived GPs to characterize disease-associated cell states across conditions and cell types, focusing on lymphoid programs that capture T cell-linked variation in AD and psoriasis (*88*). To this end, we defined disease-state metrics of dominance and specificity based on GP module scores, quantifying differences in GP activity between diseased and healthy skin (**Methods**). Applying this framework across all cell types and within lymphoid cells, including T cells, B cells, plasma and NK cells, we identified GPs with altered activity in AD and psoriasis relative to healthy skin (**Supplementary fig. S7c,d**). Given the central role of T cells in inflammatory skin disease (*89*) and evidence of persistent dysregulation skin memory T cells, including CD4+ tissue resident memory (TRM) populations in psoriasis (*90*) and broader TRM involvement across cutaneous pathophysiology (*91*), we prioritized lymphoid programs and evaluated them within skin memory T cell compartments.

To support that Tripso-derived programs reflect coordinated gene activity detectable at tissue level, we first sought orthogonal validation using matched spatial proteomics. As an illustrative example among the prioritized lymphoid programs, GP25 contains genes with epithelial and inflammatory associations (e.g. *TP63* and *CDH1*) and showed concordant spatial co-localization of representative markers at both the RNA and protein levels (**Supplementary fig. S8a**), indicating that Tripso captures diverse disease-relevant programs beyond a single cell type. We also noted that several GPs recurred in similar anatomical compartments such as epidermis- or dermis-associated regions, possibly reflecting overlapping or composite tissue states. We therefore prioritized programs with the most disease-selective profiles using the dominance and specificity metrics defined above.

Within the prioritized lymphoid programs, GP23 emerged as one of the few programs with a predominantly AD-specific profile, exhibiting high dominance and specificity for AD and low activity in healthy skin (**Supplementary fig. S7c,d**). Because regulatory T cell programs are central to maintaining cutaneous immune homeostasis and are frequently perturbed in autoimmune skin disease (*92*), we tested whether GP23 was preferentially expressed in T cell subsets with regulatory features, including IL13+ TRM cells (present in AD and healthy skin), IL17+ TRM cells (detected in Psoriasis), Type-1 regulatory (TR1), and regulatory T cells (Treg). Using pseudobulk comparisons, in which GP23 was averaged per sample within each regulatory T cell subset, we found that GP23 activity was higher in AD than in healthy skin (p < 0.05, Welch’s t-test), with no significant change in psoriasis **(Supplementary fig. S7e**). Because IL13+ TRM cells showed particularly clear separation driven by GP23 (**Supplementary fig. S7d**), we focused on this subset and confirmed the same pattern in TRM-only subsets, with GP23 significantly higher (p < 0.05, Welch’s t test) in AD than healthy skin (**Fig. 5d**). To interpret the biological basis of this signal, we next characterized the genes comprising GP23 and their differential activity across AD lymphoid subsets.

To characterize GP23 at the gene level, we focused on genes assigned to GP23 and asked which of these received higher attention in AD T cells, and in IL13+ TRM cells relative to other T cells. We tested for differential attention using the Tripso gene-discovery attention matrix (**Methods**). In the gene expression data, these genes were upregulated in AD T cells compared to psoriasis and healthy skin (**Fig. 5e, Supplementary fig. S7f**), and collectively suggest that GP23 represents a transcriptional program enriched for inflammatory signaling, metabolic adaptation, and vesicle trafficking. Indeed, these genes included inflammatory mediators (e.g., *PYCARD* (*93*), *TNFSF12* (*85*)), regulators of innate-like T cell states (*ZBTB16* (*94*), *NME1* (*95*), *FIBP* (*96*)), mitochondrial and fatty acid oxidation components (e.g. *TMEM14C* (*97*)*, COMT* (*98*)*, TSPO* (*99*)), metabolic remodelling (e.g. *FH* (*100*)) and components involved in membrane remodeling and protein turnover (*SH3GLB2* (*101*), *CUEDC2* (*102*), *PIN1* (*103*)). Many GP23 genes were not captured by existing immune pathway annotations (**Fig. 5b,f**). Notably, GP23’s closest matches in MSigDB were an immunologic signature related to *in vitro* CD4 T cell activation and a TREM2+ myeloid signature, yet overlap with these sets was limited and the majority of GP23 genes lay outside them, suggesting GP23 may represent a previously unannotated transcriptional program. Collectively, these results illustrate how moving beyond canonical cell-type markers and established GP databases enables a more functionally informed characterization of disease-associated cell states.

Given that skin-resident memory T cells occupy and are maintained within localized epidermal and appendage-associated niches supported by local cytokines (*104*), we asked whether GP23 exhibited a corresponding spatial signature in tissue. In a representative lesional AD relapse section collected 8 weeks after treatment (Dupilumab) withdrawal, hematoxylin and eosin staining revealed dense lymphocytic aggregates (**Fig. 5g**). In the same region, GP23 expression was spatially enriched in discrete immune-dense niches adjacent to sebaceous glands and within inflamed epidermis (**Fig. 5h,i**). This pattern was reproduced in an independent cohort of Xenium sections of AD lesional skin (n= 3 patients) (**Fig. 5j; Supplementary fig. S9a–d**), where GP23-high regions were consistently observed in sebaceous and epidermis-associated inflammatory areas in close proximity to T cell aggregates. Across 60 Xenium section from 19 patients (15 AD and 4 psoriasis; 44 AD and 16 psoriasis sections), expression of GP23 constituent genes was highest in sebaceous gland and inflammatory epidermis niches, with clearer distinction in immune-associated sebaceous and plasma cell–rich niches (**Fig. 5f**). This pattern was consistent across patients, sections and recurred across distinct inflammatory niche categories, supporting reproducible, niche-restricted GP23 expression in tissue.

To show the link between GP23 expression and immune infiltration, we analyzed matched spatial proteomics from an AD follow-up section (week 12 after treatment), a stage associated with ongoing tissue remodeling and persistence of resident immune populations. We found that GP23-high regions frequently coincided with areas of high CD45 (PTPRC) protein signal, a pan-leukocyte marker (**Fig. 5j,k**), consistent with immune enrichment in GP23-high niches. We used epithelial markers in the same panel (e.g., E-cadherin) to delineate keratinocyte- and ductal-associated compartments, enabling interpretation of GP23-high regions in relation to both immune and epithelial tissue structures (**Supplementary fig S8a**). Across multiple sections, GP23-high sebaceous regions contained epithelial cells (keratinocytes and sebaceous/ductal compartments) which lay in close proximity to T helper cell aggregates (**Supplementary fig. S8b-c**), suggesting that GP23 reflects a niche-associated tissue state linked to immune proximity, rather than a simple proxy for immune cell abundance. Consistent with this, defining T cell hotspots based on local T cell density (**Methods**) revealed that GP23 activity decreased with increasing distance from the nearest hotspot (**Supplementary fig. S10a-e**). Together, these results link the GP23 signal observed in dissociated T cells to a niche-restricted program enriched in sebaceous- and epidermis-adjacent regions near T cell aggregates, consistent with TRM cells adopting a distinct metabolic state in lipid-rich tissue environments to support their persistence (*105*).

In AD, GP23 module scores in IL13+ TRM cells were lowest in healthy control skin, increased in untreated lesional skin, and remained elevated in relapsed skin sampled 8 weeks after Dupilumab withdrawal (**Fig. 5m**), pointing to enrichment of GP23-high niches in active and recurrent disease. Notably, GP23 module scores were higher in relapsed skin than untreated lesional samples, suggesting that disease recurrence is associated with expansion or increased abundance of GP23-high niches rather than simple reversion to the prior lesional state. Although GP23 was not globally elevated in psoriasis compared to healthy skin (**Fig. 5d**), GP23 activity was detected in resolved psoriasis samples (**Supplementary fig. S11a-c**), indicating that GP23-associated niches can persist beyond clinical resolution in distinct inflammatory contexts. This pattern aligns with the capacity of tissue-resident memory T cells to endure after lesion clearance and contribute to recurrent inflammation. Together, these results demonstrate that clustering Tripso attention profiles yields interpretable GPs with reproducible spatial organization in inflammatory skin.

## Discussion

Single-cell atlases are increasingly comprehensive, yet many downstream applications of these data rely on compressing cell state into a single latent representation, or simplify pathways into a single score representing pathway activity. This representation makes it difficult to separate concurrent GPs, such as transcription factor regulons and signaling responses, that coexist within the same cell and vary across batch, tissues, ages, genetic backgrounds and *in vitro* conditions. Tripso addresses this gap by reframing cellular state as a composition of interpretable GP embeddings, combining the representational flexibility of transformers with biological interpretability in single-cell analysis. Across controlled perturbations and complex tissues, Tripso improved discrimination of GP activity. Applied to hematopoiesis across the lifespan, Tripso refined the characterization of cell states beyond static cell type labels. By comparing hematopoietic cells *in vivo* and *in vitro*, Tripso enabled actionable, program-level comparisons between these systems. In inflammatory skin disease, Tripso identified data-derived GPs that distinguished disease-associated immune states and showed coherent, reproducible spatial patterning when mapped back onto tissue architecture.

Tripso advances single-cell analysis by replacing scalar pathway scores with multidimensional, GP-resolved representations that capture context-dependent biological variation. Existing label-free approaches either summarize GP activity as a single score or constrain each program to a single latent dimension, limiting their ability to represent heterogeneous GP usage across cell states. In contrast, Tripso learns flexible, GP-specific embeddings without requiring cell-type labels, enabling analysis in settings where canonical identities are ambiguous or unavailable. These embeddings interoperate seamlessly with existing single-cell analysis frameworks, including differential composition analysis and optimal transport, extending their application from cell to GP-level interpretation. Together, these features establish Tripso as an advance beyond GP-centric scoring toward GP-aware representations that preserve biological complexity while remaining interpretable.

Tripso reveals biological structure can be obscured in global representations, where concurrent programs are conflated. In hematopoiesis, this is illustrated by early B-lineage cells, where prenatal and postnatal populations separate specifically within the IKZF1 GP. The observed pattern is consistent with a developmental transition from proliferative expansion in prenatal stages to increased diversification postnatally, including enhanced BCR diversification, as supported by both single-cell transcriptomic and BCR sequencing data (*63*). Notably, these populations remain indistinguishable when examined through unrelated programs, highlighting how GP-resolved embeddings provide precise, context-specific resolution of cell state.

Embedding *in vivo* and *in vitro* HSCs into Tripso GP space enabled systematic linkage of GP to stem cell states and highlighted tractable intervention points. Tripso identified SEC61 inhibition as a candidate strategy to improve immunophenotypic HSC maintenance *in vitro*. *SSR1* and *SEC61G*, upregulated in lineage-committed progenitors relative to stem-like cells, encode components of the endoplasmic reticulum (ER) translocon, which mediates co-translational protein targeting to the ER. Disruption of this process can activate the unfolded protein response (*106*), a pathway elevated in quiescent HSCs (*107*) and rapidly diminished in culture (*108*). We speculate that SEC61 inhibition may partially restore this stress-adapted state under permissive culture conditions, or preferentially impair progenitors with higher translational output, thereby enriching stem-like cells. Notably, although *SSR1* and *SEC61G* are differentially expressed and thus detectable by conventional analyses, situating them within GPs provides a principled framework for prioritization and interpretation. This example illustrates the broader potential of GP-resolved representations to systematically dissect complex cellular states, inform the optimization of therapeutically relevant protocols and guide hypothesis generation across biological contexts. This is analogous to work carried out in Genes2Genes, where trajectory alignment of T cell differentiation *in vivo* and *in vitro* identified TNF signalling as a target to improve in vitro differentiation (*109*). However, unlike Genes2Genes, which performs alignment at the level of individual genes, Tripso models GPs as a primary representation, aggregating signals across genes and reducing sensitivity to noise or dropout, while mitigating reliance on explicit pseudotime calculation.

In inflammatory skin disease, applying Tripso to large-scale single-cell data integrated with matched spatial transcriptomic and proteomic measurements enabled contextualization of data-derived GPs within tissue architecture. Tripso-derived GPs exhibited limited overlap with curated gene-set databases, delineated disease-associated immune cell states, and displayed niche-restricted expression within tissue. Notably, IL13+ TRM cells have been reported to localize adjacent to sebaceous glands (*86*), a lipid-rich tissue compartment, and TRM persistence depends on exogenous lipid uptake and metabolism (*105*). These features suggest that AD IL13+ TRM cells undergo mitochondrial and metabolic adaptations to persist within such niches, thereby supporting chronic inflammation and disease recurrence. The persistence of GP23-associated niches in resolved (lesion-cleared) skin further supports the idea that spatially restricted immune programs can endure beyond clinically visible inflammation and contribute to disease recurrence (*110*). The presence of skin-resident memory T cells in relapse or lesion-clear samples is associated with GP23 high niches, which can act as a supportive transcriptional program to sustain their survival. Together, these results highlight how attention-based GP discovery can enable data-driven identification of spatially organized transcriptional states across conditions.

Tripso does have several limitations. The framework relies on a predefined set of GPs, which enhances interpretability but constrains analysis to the scope and quality of the input program definitions. GPs absent from this set cannot be directly interrogated, although the GP discovery module provides a mechanism to identify dataset-specific programs and partially expand coverage. Tripso further assigns a dedicated transformer block to each GP, enforcing parameter independence across programs. While this architectural choice limits scalability and necessitates prior curation of GPs, it is well suited to hypothesis-driven analyses in which the biological processes of interest (such as signaling pathways or transcription factor regulons) are known a priori and require interpretable, GP-specific representations. Finally, in its current implementation, Tripso operates on transcriptomic data alone, which may limit resolution for programs governed primarily by chromatin accessibility, protein abundance, or spatial context, motivating future extensions of the framework to multimodal single-cell data.

Looking ahead, Tripso defines a GP-level representation of cellular state that provides a foundation for future methodological extensions. Although applied here to transcriptomic data, the framework could be extended to additional modalities by modeling GPs across paired measurements, such as chromatin accessibility. In spatial transcriptomics, one possible extension would incorporate spatial proximity directly into the representation learning objective, for example, by encouraging cells that are physically adjacent to share similar GP embeddings, thereby enabling explicit modeling of niche-level organization rather than post hoc spatial validation. While we focus here on human data, recent work suggests that GPs may constitute an evolutionary unit shaping cellular phenotypic and functional novelty (*111*), making GP-level representations a natural substrate for cross-species comparison even when individual genes or cell types are not directly conserved. Such extensions could leverage protein-sequence–based embeddings to initialize gene representations, as demonstrated in cross-species foundation models such as UCE (*12*). Finally, the growing availability of large-scale perturbation screens with functional readouts, such as Perturb-FISH, which pairs genetic perturbation with imaging-based functional readouts (*112*), will enable the construction of increasingly high-quality, functionally grounded GP libraries for comparison across tissues, conditions, and disease states.

More broadly, Tripso reframes how cellular states are compared across biological contexts by moving beyond forced integration and towards GP-aware comparison. This shift enables researchers to identify which GPs are conserved, which diverge, and which represent tractable targets for intervention. By enabling interpretable, GP-focused analysis at scale, Tripso enables new ways in which single-cell data can be used to generate hypotheses, guide experimental design, and translate molecular insight into actionable biological and therapeutic advances.

## Data Availability

We provide our newly generated single-cell, CITE-seq and spatial data as an interactive resource (https://cellatlas.io/tripso), with Tripso GP embeddings and novel GP module scores. Raw sequencing data for the hematopoietic MNC atlas are publicly available via GEO or ArrayExpresss with accession numbers listed in **Supplementary table t7**. All other data will be made available upon publication.

## Code Availability

Tripso is available as a Python package, maintained at https://github.com/Lotfollahi-lab/tripso. All code to reproduce benchmarking and analyses is available at https://github.com/Lotfollahi-lab/tripso_reproducibility. We provide documentation including tutorials and a user guide at https://tripso-docs.cog.sanger.ac.uk/.

## Acknowledgements

We thank Aidan Maartens for editorial review of the manuscript. We are further grateful to all members of the Vento-Tormo, Haniffa, Göttgens, and Lotfollahi teams for their constructive discussions and feedback throughout this project. We acknowledge core funding from Wellcome (WT220540/Z/20/A). This project was conceived and funded by Open Targets, under OTAR 3090. This research was funded in whole, or in part, by the Wellcome Trust [203151/Z/16/Z, 203151/A/16/Z, 226795/Z/22/Z] and the UKRI Medical Research Council [MC_PC_17230]. We thank the teams of Anthony Nolan Tissue Bank and the Cambridge Blood and Stem Cell Biobank for provision of cord blood samples. M.H. is funded by Wellcome (WT107931/Z/15/Z) and the CIFAR MacMillan Multiscale Human program. EL was funded by a Sir Henry Dale fellowship from Wellcome/Royal Society (107630/Z/15/Z). Research in EL’s laboratory was/is funded by Biotechnology and Biological Sciences Research Council (BB/P002293/1) and by Wellcome (215116/Z/18/Z and 309075/Z/24/Z). N.M. was supported by the Japan Society for the Promotion of Science Short-term Postdoctoral Fellowship and a Deutsche Forschungsgemeinschaft (DFG) Research Fellowship (ME 5209/1-1). T.I. was supported by the Funai Foundation for Information Technology and the Honjo International Scholarship Foundation. Work in the Gottgens laboratory was supported by Wellcome Trust [206328/Z/17/Z, 215116/Z/18/Z, 221052/C/20/Z], Medical Research Council [MR/W031663/1 and MR/V005502/1], Blood Cancer United [7035-24], Blood Cancer UK [18002] and Aging Biology Foundation. LPC is funded by Wellcome Trust [318995/Z/24/Z]. S.W. is funded by the Royal Society (CDF/R1/241008), and supported by the NIHR and Newcastle Biomedical Research Centre. D.J is supported by a Wellcome Trust Accelerator Award (314710/Z/24/Z) and the NIHR Biomedical Research Centre at Great Ormond Street Hospital for Children NHS Foundation Trust and University College London. L.S. is supported by a Wellcome Clinical Research Fellowship PhD student grant and Trinity Internal Graduate Studentship. M.H. is funded by Wellcome (grant WT107931/Z/15/Z) and the NIHR Newcastle Biomedical Research Centre. SKM is funded by a National Institute for Health and Care Research (NIHR) Advanced Fellowship (NIHR302258). For the purpose of Open Access, the authors have applied a CC BY public copyright licence to any Author Accepted Manuscript version arising from this submission. Fig. 1 (all panels), and Figures 2a, 2g, 3a, 4a, 4e and 5a were created using Biorender.

## Author Contributions

ML, NKW, MH, BG and RVT conceptualized the study. AV, MM conceived and developed the method with feedback from ML, RVT, and implementation help from CS, KCHL, HA. ML, BG, MH, RVT, NKW, MM, TI, AV, CL designed experiments. MM, TI, CL analyzed the data, with help and feedback from AV, NKW, EW, VS, AM, CRS, LS, JYWL, DJJ, SM, CS, RVB. LPC carried out the expansion culture experiments. MP, AH, DH, DBL, provided software development support. Blood related CITE-seq datasets were generated by NM, ES, DI, and NKW. AR, IR and EL provided resources and supervised the generation of the blood CITE-Seq datasets. Computational analysis and QC of CITE-Seq datasets was performed by TI, MQL, SW, IG, MSV, QL and RH. ARF, BR, HC, performed Xenium ST experiments. ARF and KR performed the Akoya Phenocycler validation. ML, NKW, MH, RVT, BG supervised the project. All authors read and approved the manuscript.

## Methods

### Overview

Tripso is a self-supervised framework for learning interpretable single-cell representations through a GP lens. Tripso operates in three stages: (i) learning gene-level embeddings, (ii) learning embeddings for each curated GP independently, and (iii) combining GP embeddings to construct a global representation of each cell.

### Tokenization and gene encoding

We follow the rank-based tokenization introduced by Genformer (*8*), restricting inputs to genes that belong either to curated GPs or to a set of highly variable genes (HVGs). HVGs were obtained either directly from the original publication or computed using Scanpy (*1*) with the seurat_v3 method; when datasets combined multiple studies, study was used as the batch key. The maximum sequence length for a given cell is the union of GP genes and HVG. Including HVGs alongside GP member genes allows the model to retain additional sources of transcriptional variability not explicitly captured by curated GPs, while restricting to these genes mitigates the computational cost of self-attention, which scales quadratically with sequence length.

Each gene is represented as a token and passed through a transformer encoder (hereafter referred to as the gene encoder), which produces gene-level embeddings. To leverage information learned from large-scale pretraining, the gene embedding tokens matrix is initialized as a look-up table using pretrained gene embeddings from Geneformer (4096 12L 96M model (*113*), accessed May 2025). The gene encoder is trained using a masked language modeling objective over gene tokens, enabling it to learn context-dependent expression relationships. If **M**_*i*,*g*_ denotes the number of masked gene tokens in the full sequence for cell *i*, and 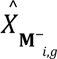 denotes the corresponding unmasked gene embeddings, then the loss for the gene encoder is then defined as:

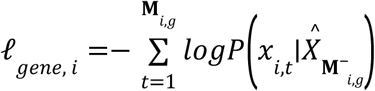

### GP representation learning

We define a collection of curated GPs, where *J* is the total number of curated GPs and each GP consists of a set of biologically related genes (see GP curation below). While the number and size of GPs are flexible, computational cost scales with the number of GP blocks, as each is parameterized independently.

Each GP is modeled independently using a transformer encoder with its own parameters. Attention is implemented using the PyTorch FlashAttention backend for improved memory efficiency and computational speed. For a given cell, the gene-level embeddings produced by the gene encoder are filtered to retain those corresponding to genes in GP *j*. A learnable GP-specific CLS token is prepended to this sequence, and the resulting embeddings are passed through the GP transformer. The goal is to learn a set of GP-level embeddings, where the GP CLS summarizes the activity of GP *j* in cell *i*.

GP transformers are trained using a bidirectional masking objective analogous to BERT (*6*). Gene tokens within each GP are randomly masked with probability *p* (0.25 by default), the CLS is never masked. Among masked tokens, 10% are replaced with random tokens and 10% are left unchanged, which improves generalization. The model is trained to predict the identity of masked gene tokens based on the remaining unmasked genes within the same GP. For GP *j*, let **M**_*i*,*j*_ denote the number of masked gene tokens within GP *j* for cell *i*, and let 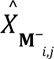 denote the set of unmasked gene embeddings within GP *j* for cell *i*. The masking loss is defined as

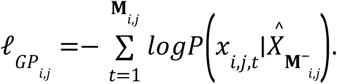

The total GP loss is defined as the sum of masking losses across all GPs.

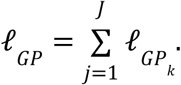

We refer to the collection of GP transformers and the gene encoder as the base model. The loss for the base model is thus

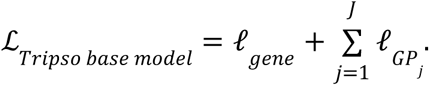

### Global cell representation

Using the learned GP embeddings, we construct a global representation for each cell. For each cell, we assemble a sequence of GP CLS tokens (ordered based on the proportion of GP genes expressed in cell *i*) and prepend a learnable cell-level CLS token. This sequence is passed through a transformer encoder to produce the global cell embedding.

To encourage the cell embedding to capture biologically meaningful variation, we train a decoder to reconstruct original gene expression. Raw counts are filtered to the same gene set used for tokenization (union of GP member genes and HVGs). The decoder consists of a two-layer multilayer perceptron with Euclidean normalization, followed by a reconstruction loss based on the negative binomial (NB) distribution, as implemented in scVI (*2*).

### Model training

Model training proceeds in three phases to promote stability and effective learning across hierarchical levels. First, the gene encoder is trained using a gene-level masking objective for 10 epochs. Training continues for an additional 10 epochs using a combined objective that includes both gene-level masking and GP-level gene masking. Finally, the global cell model is trained for 5 epochs using only the gene expression reconstruction loss. During this phase, the gene encoder and GP encoders are frozen, and only parameters involved in constructing the global cell embedding and decoder are updated.

### Discovering novel gene programs

We implemented a GP discovery module to identify biologically meaningful gene modules from a user-defined collection of input genes, such as HVGs or predefined panels from other modalities (e.g., spatial transcriptomics).

After training the base model (gene encoder and curated GP transformers), we initialize an additional transformer block dedicated to GP discovery. This block takes the selected gene set as input. Unlike the base model, masked gene modeling is not applied at this stage. Instead, the discovery transformer learns a CLS token which is passed to the global cell module and trained using the gene expression reconstruction objective. During this phase, only the discovery transformer is updated, while the pretrained gene encoder and curated GP transformers remain fixed. This design allows the model to learn gene-gene relationships within the selected gene set while leveraging contextual information captured during base model training.

After training, the GP discovery module produces CLS-to-gene attention weights that support two complementary analyses. First, in order to rank genes associated with a specific cell population or condition (e.g., cell type, disease status), we compute the mean CLS-to-gene attention within that group and compare it to the mean attention across all other cells, ranking genes by their differential attention. Second, GPs are identified by clustering genes with similar CLS-to-gene attention patterns across cells. GPs were derived from this CLS-to-gene attention matrix using two complementary strategies described below.

#### Lineage-conditioned GPs

To identify GPs within lineages, we analyzed CLS-to-gene attention patterns within specific cell populations (e.g., cell types). For each population, we restricted analysis to the relevant subset of cells, excluding populations with fewer than 20 cells.

Within each population, we computed a gene-gene similarity matrix based on the absolute Pearson correlation of gene attention scores across cells. Genes were then clustered using spectral clustering on this similarity matrix, with each cluster defining a candidate GP. To determine the number of clusters (K), we evaluated a range of candidate values and selected the K that maximized the average silhouette score, thereby identifying the most well-separated clustering solution.

Each resulting gene cluster defines a lineage-conditioned GP. In the main text, GPs were subsequently assigned unified identifiers (GP_1_, GP_2_, …, GP_N_).

#### Multilineage gene programs

To identify GPs that reflect coordinated activity across multiple lineages, rather than those restricted to a single lineage, we implemented a complementary strategy inspired by the frequency matrix framework described in CoVarNet (*114*). In this section, we included multiple cell types from distinct lineages as input, rather than analyzing each cell type separately.

In order to account for possibly different distributions of attention scores between different cell populations, we used a rank-based representation to identify genes that are consistently prioritized across cells. For each cell, we defined salient genes as the genes whose CLS-to-gene attention scores fell within the top 3% of that cell’s distribution, yielding a sparse binary matrix that indicates whether each gene is highly attended to in a given cell.

Next, we defined cellular contexts as the Cartesian product of cell-type annotation and sample of origin (i.e., each context corresponds to a specific cell type-sample pair). Contexts containing fewer than 20 cells were excluded. Within each retained context, for every gene, we calculated the proportion of cells in which that gene was classed as salient. This produces a gene-by-context matrix F that quantifies how frequently each gene is strongly attended to in each context.

Finally, we computed pairwise Pearson correlations between genes based on their context-frequency profiles encoded in the rows of F, yielding a gene-gene similarity matrix. We then clustered genes using spectral clustering on the absolute correlation matrix. As in the lineage-specific setting, the number of clusters K was selected by maximizing the silhouette score over a range of candidate values. Each resulting cluster defines a multilineage GP. GP were retained if their size fell within a predefined range (25–250 genes).

For both lineage-conditioned GPs and multilineage GPs, we explored their expression in original gene expression space using Scanpy module score (*1*).

### Metrics

#### Quantifying gene contributions to GP representations

To quantify the contribution of individual genes to GP representations, we compute a gene importance score for each cell as the cosine similarity between the gene embedding and the corresponding GP CLS within that cell. Higher cosine similarity indicates a stronger contribution of the gene to the GP representation. To identify genes that are differentially important between two classes, we first remove genes that are missing in more than a user-defined fraction of cells. For remaining genes, if a gene is absent in a given cell, its importance score is set to zero, reflecting its lack of contribution. Differential importance is computed as the difference in mean cosine similarity between classes. Statistical significance is assessed using a t-test, and p-values are corrected for multiple testing using the Benjamini-Hochberg correction.

#### Quantifying GP contributions to cell representations

To assess the contribution of each GP to the global cell representation, we perform post hoc feature ablation. After training the Tripso cell decoder, model parameters are frozen. For each GP, we set its embedding to zero and recompute the cell embedding.

GP importance for a given cell is defined as follows:

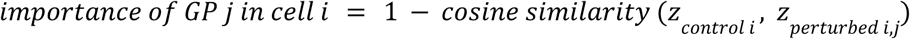

where *z*_*control i*_ is the original representation for cell *i*, and *z*_*perturbed i,j*_ is the output cell representation when the GP *j* embedding is set to 0. This formulation ensures that larger values correspond to greater influence of the GP on the cell representation.

To test for differential GP importance between classes, we construct an AnnData (*115*) object of shape (N, J) where J is the number of GP and each entry represents the importance score of a GP in a given cell. Differential importance is computed using the t-test implemented in Scanpy’s rank_genes_groups (*1*), and GPs are ranked by log-fold change.

#### Linear probing

To evaluate how well learned embeddings discriminate between biological classes, we train logistic regression classifiers (implemented in scikit learn (*116*)) on GP embeddings to predict labels of interest. We use weighted logistic regression to account for class imbalance and report class-specific F1 scores.

#### Changes in GP importance as a function of differentiation

To identify GPs whose importance varies systematically with differentiation, we fit a separate generalized linear model (GLM) for each GP. The response variable is the GP importance score for each cell, and the predictor is a numeric encoding of cell type ordered by differentiation state. Models are fit using a Gaussian family with identity link via the statsmodels library (*117*). GPs are ranked according to the magnitude of the regression coefficient associated with differentiation.

#### Optimal transport for mapping across conditions

To infer correspondences between cells across experimental conditions, we apply optimal transport (OT) to the learned cell embeddings. We use the framework implemented in the OTT-JAX library (*118*), following the approach of moscot (*119*). OT identifies a coupling between two cell populations that minimizes transport cost, defined as squared Euclidean distance between embeddings. Because exact OT is computationally expensive, we use entropically regularized OT solved via the Sinkhorn algorithm, which improves numerical stability and scalability (*120*). We set the regularization strength to 10^-3^. To account for differences in population composition across conditions, we employ unbalanced OT, which relaxes marginal constraints using Kullback–Leibler divergence penalties (*121*). The scaling parameters controlling the marginal relaxation were set to 0.999 for both distributions. Prior to computing OT, embeddings were reduced to their first 50 principal components. OT couplings were then computed in the reduced space. To mitigate the effects of severe class imbalance, we subsampled cells to achieve approximate balance between conditions. For visualization, we identified representative cells closest to cluster centroids and examined OT mappings between these points.

To quantify population-level similarity between conditions, we computed the Sinkhorn divergence (*122*) as implemented in OTT-JAX (*118*). Sinkhorn divergence corrects for entropic bias by subtracting self-similarity terms, and provides a stable, positive-definite measure of distributional similarity.

#### Quantifying condition-level GP activity, dominance and specificity

To compare GP activity across conditions while accounting for differences in cell-type composition, we define the following metrics of GP activity.

##### Condition-level GP activity

We first defined GP activity based on module score (*1*) in the gene expression data. For each GP *j*, we compute a per-cell module score. We then calculate the mean GP activity µ_*j*,*d*,*t*_ within each condition *d* and cell type *t* (in our main analysis, condition refers to healthy, atopic dermatitis or psoriasis but condition can represent any relevant metadata). To obtain a single condition-level estimate that accounts for variation in cell-type composition, we compute µ_*j*,*d*_, the average of µ_*j*,*d*,*t*_ weighted by the number of cells from each cell type.

##### Condition Dominance

To quantify how strongly a GP is associated with each condition relative to others, we defined condition dominance as the normalized share of total GP expression attributable to that condition:

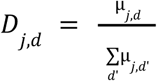

where d’ represents the different condition values in the dataset. By construction, *D*_*j*,*d*_ ∈ [0, 1], and the dominance values for a given program sum to 1 across conditions. Dominance therefore captures programs whose overall activity is predominantly attributable to a given condition.

##### Condition Specificity

To quantify condition-specific shifts in GP activity relative to a reference state (eg: healthy), we defined specificity as the positive difference in condition-level activity:

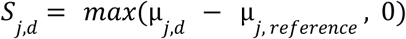

This metric captures the magnitude of program upregulation in a given condition relative to the reference, with negative differences truncated to zero.

These metrics were computed either across all cells or within specified cellular compartments (e.g., lymphoid populations), using the same definitions.

### Novelty and Reconstructability of Tripso GPs against curated gene set resources

To assess whether Tripso-derived GPs correspond to previously published pathways and gene expression signatures, we compared each GP to public gene set collections using two complementary approaches: (i) best single-set similarity and (ii) multi-set reconstruction. In this section, we will use the term GP to refer to Tripso-derived GPs and gene sets to refer to public gene sets.

We downloaded human gene sets from the Molecular Signatures Database (MSigDB, v2025.1). We included the Hallmark collection (H, 50 curated gene-sets) which spans broad transcriptional states, Canonical Pathways (C2, 4115 curated gene-sets) to represent signaling and metabolic pathways relevant to inflammatory tissue remodeling, Immune Perturbations (C7, 4872 curated gene-sets) that reflect immune activation and perturbation signatures which may be pertinent to T cell-driven disease, Cell Type Signatures (C8, 866 curated gene-sets) which could help assess whether inferred GPs reflect cell-type markers. For comparisons to pretrained programs used in the base model, we also included PROGENy pathway gene sets (14 signaling pathways). Because Tripso GPs were derived from the 5,000 genes included in the Xenium panel, all comparisons were restricted to these genes, and public gene sets were filtered accordingly prior to analysis.

We then computed the Jaccard similarity between each Tripso GP and each public gene set. For each Tripso GP, we recorded the maximum Jaccard similarity within each collection and the identity of the corresponding best-matching set. We also recorded the maximum similarity across the combined MSigDB collections.

To test whether a GP could be approximated by combining multiple gene sets, we implemented a greedy reconstruction procedure that penalizes false positives. Starting from an empty set, we iteratively added the gene set that maximized coverage of the GP while penalizing inclusion of off-target genes (penalty weight = 0.2). Only gene sets sharing at least two genes with the GP were considered. Iteration continued until either 75% of the GP genes were covered or a maximum of 60 gene sets had been selected. Reconstruction performance was summarized by: K₇₅ the number of gene sets required to reach 75% GP coverage (capped at 60), and the Jaccard similarity between the GP and the union of selected gene sets. GPs that did not reach 75% coverage within 60 sets were classified as not reconstructable under this criterion.

### Datasets

For each datasets, we randomly split 80%, 10%, 10% of the data into training, validation and test sets. For benchmarking experiments, we use the same splits across methods, and report all results on the test set. For the hematopoiesis and skin results sections, results are shown on the combined splits in order to maximize available data for biological discovery.

#### Perturbation dataset

Gene expression data was downloaded from https://zenodo.org/records/10520190 (*30*). The original dataset contains 1.6 million cells representing stimulations for five signalling pathways: IFNβ, IFNγ, TNFα, insulin and TGFβ. In our benchmark experiment, we aim to leverage these stimulations to test different methods’ ability to separate cells based on ground truth GP activity. However, inflammatory pathways such as IFNβ, IFNγ, and TNFα may produce similar responses, and it is not necessary that increased discrimination of these classes indicates better reflection of true biological processes. Therefore, to simplify interpretation, we restrict our analysis to single inflammatory stimulation, TNFα. Furthermore, insulin simulation induced heterogeneous response across different cell lines and the authors’ analysis did not identify a consensus set of insulin response genes(*30*). Therefore, for the purposes of benchmarking, we also excluded insulin-stimulated cells and focused on TGFβ and TNFα stimulated cells.

To determine the genes which induce the strongest perturbation effect in each pathway, we use the cell-level Mixscale score, a scalar value reflecting the strength of perturbation response developed by the original authors. We computed the average gene-level Mixscale, and ranked the genes by most- to least- pronounced average effect.

##### GP curation

We use TGFβ and TNFα response signatures provided by PROGENy(*31*) as GP inputs to Tripso, focusing on the 200 genes with the smallest p-values associated with pathway response (**Supplementary table t6**). We include up to 200 genes with significant associations, leveraging transformer models’ ability to flexibly attend to distinct gene subsets in individual cells, enabling model-driven refinement of GP genes.

### Human Endometrial Cell Atlas

Gene expression data was downloaded from https://www.reproductivecellatlas.org/endometrium_reference.html (*32*). Raw counts were used as tokenization inputs without further preprocessing.

#### GP curation

As above, we use the transcriptional response signatures provided by PROGENY(*31*), selecting the 200 genes with the smallest pathway-associated p-values (**Supplementary table t6**). The TRAIL GP was excluded because its target genes showed low expression, with an average of <10 genes with non-zero expression per cell.

### Hematopoiesis dataset

#### Tissue acquisition

Human developmental tissues (YS, FL and fetal BM) were obtained from the Human Developmental Biology Resource (HDBR), following elective termination of pregnancy, with written informed consent and approval from the Newcastle and North Tyneside NHS Health Authority Joint Ethics Committee (18/NE/0290 and 08/H0906/21+5). The HDBR is regulated by the UK Human Tissue Authority (HTA; https://www.hta.gov.uk/) and operates in accordance with the relevant HTA Codes of Practice. Cord blood samples were obtained from Cambridge Blood Stem Cell Biobank, in accordance with regulated procedures approved by the relevant Research and Ethics Committees (18/EE/0199). Cord blood samples were purchased from Anthony Nolan (REC ref 20/EM/0028) for in vitro culture experiments. Aged adult tissues were obtained from Newcastle Hospitals NHS Foundation Trust, following hip replacement surgery, with written and informed consent and approval from the Newcastle and North Tyneside NHS Health Authority Joint Ethics Committee (19/LO/0389).

#### Tissue processing

Adherent material was removed, and the fetal femur was cut into small pieces before grinding with a pestle and mortar. Flow buffer (PBS containing 5% (v/v) FBS and 2 mM EDTA) was added to reduce clumping. The suspension was filtered with a 70-μm filter and red cells were lysed with 1× RBC lysis buffer (eBioscience) according to the manufacturer’s instructions.

#### CITE-seq experiments

Cryopreserved YS (n = 2; 6-7 PCW), FL (n = 9; 6-17 PCW), fetal BM (n = 4; 14-17 PCW), cord blood (n = 4) and adult BM cells (n = 9; 29-91 years old) (**Supplementary table t7**) were thawed on the day of experiment and added to pre-warmed RF-10 (RPMI (Sigma-Aldrich) supplemented with 10% (v/v) heat-inactivated FBS (Gibco), 100 U ml^−1^ penicillin (Sigma-Aldrich), 0.1 mg ml^−1^ streptomycin (Sigma-Aldrich), and 2 mM l-glutamine (Sigma-Aldrich). Cells were manually counted and pooled if cell numbers were low (pools noted in the sequencing index columns; **Supplementary table t7**). Cells were incubated with Fc receptor blocking reagent (BioLegend) for 10 min in the dark and on ice. During the incubation, the CITE-seq antibody cocktail vial was centrifuged at 14,000g for 1 min then reconstituted with flow buffer. The vial was incubated for 5 min at room temperature then centrifuged at 14,000g for 10 min at 4 °C. The CITE-seq antibody cocktail (**Supplementary table t8**) was then added to the cells along with a competition antibody mix (**Supplementary table t9**) and incubated for 30 min in the dark and on ice. The stained cells were then washed with flow buffer before resuspension in flow buffer supplemented with 50 μg ml^−1^ 7-AAD (Thermo Fisher Scientific).

Live, single cells were sorted by FACS into 500 μl PBS in pre-chilled FACS tubes coated with FBS until the sample was exhausted. Sorted cells were then centrifuged at 500g for 5 min before manual counting. Cells were then submitted to the CRUK CI Genomics Core Facility for 10x Chromium loading, library preparation and sequencing. Single cell 3′ v3 (10x Genomics) kits were used and gene expression and cell-surface protein libraries were generated as per the manufacturer’s protocols. Libraries were sequenced using an Illumina Novaseq platform. The CD34^+^ YS, FL, fetal BM, CB and PBM samples have been published as part of our previous studies(*36*, *37*, *123*), as well as the combined CD34^+^ atlas (*40*).

#### Alignment and preprocessing of CITE-seq data

CITE-seq transcriptomic data were aligned to the GRCh38 genome and quantified using the Cell Ranger pipeline (v4.0.0). Cell-associated barcodes and background-associated barcodes were determined using the EmptyDrops (*124*) method implemented in the Cell Ranger pipeline, and the background-associated barcodes were excluded. Pooled samples were demultiplexed with Souporcell (*125*) and Vireo (*126*), where individual donor-derived cells were first clustered using Souporcell and donor information was matched using Vireo based on paired SNP array data (UK Biobank Axiom Array, Applied Biosystems, performed by Cambridge Genomic Services, University of Cambridge) obtained for each pooled donor. Subsequent data analysis was performed using Scanpy (*1*). Multiplets were removed using Scrublet (*127*) based on the threshold of the median plus three times the median absolute deviation scrublet score, as previously described (*35*). Cell libraries with less than 2,000 total unique molecular identifier (UMI) counts or with mitochondrial gene expression exceeding 10% of UMI counts were removed.

Cell surface protein data were quantified using CITE-seq-Count (*128*) (v1.4.3) with the option ‘-cells 200000’ to return sufficient background droplets for the subsequent denoising analysis. Cells with more than 160 detected proteins or with total protein counts less than the median minus 0.8 times the median absolute deviation were removed. Data for the remaining 82,035 cells were denoised and normalized using DSB(*129*). The DSB-normalized protein expression matrix was used to compute 50 principal components (PCs), and the PCs were adjusted for batch effects between the individual samples using Harmony(*130*). The 50 batch-corrected PCs were used to identify 12 nearest neighbors and to compute clusters using the Scanpy functions ‘pp.neighbors’ and ‘tl.leiden’, and the following multiplet clusters were removed: HSPC/myeloid/B-cell multiplets (CD34^+^ CD15^+^ CD19^+^), B/myeloid doublets (CD19^+^ CD33^+^), erythroblast/monocyte doublets (CD235a^+^ CD14^+^), and T/myeloid doublets (CD3^+^ CD33^+^ CD123^+^).

### Dimensional reduction and annotation

The gene expression count matrix for the remaining 80,794 cells was log-normalized, and 2,000 highly variable genes were identified. Cell cycle-associated genes (*131*) were then removed and the expression values of the remaining 1,908 highly variable genes were scaled and used to compute 50 PCs. Protein expression-based PCs were recomputed for the remaining 80,794 cells. The transcriptome- and protein-based PCs were separately adjusted for batch effects between the individual samples using Harmony (*130*). A multimodal neighbor graph was constructed based on the 50 transcriptome-based and 20 protein-based batch-corrected PCs using the weighted nearest neighbor (WNN) algorithm (*132*). The WNN graph was then used to compute clusters and the UMAP embedding using the Scanpy functions ‘tl.leiden’ and ‘tl.umap’, respectively. Data annotation was performed manually using marker genes and proteins identified through literature search (**Supplementary table t4**). Stromal cell populations, including *CDH5*^+^ endothelial cells (*133*), *COL1A1*^+^ *DCN*^+^ fibroblasts (*134*), *ALB*^+^ *AFP*^+^ hepatocytes (*135*), and *ACP5*^+^ *CTSK*^+^ osteoclasts (*136*) were also identified and excluded from our hematopoietic landscape. For the remaining 79,583 cells, the final UMAP embedding was recomputed using the ‘tl.umap’ function.

#### Data curation

Our research questions were two-fold: first, to explore how GP activity changes across hematopoietic tissues and life stages, and second, to assess the extent to which patterns of GP activity are conserved *in vitro*. To learn generalizable GP representations, we assembled a training corpus of 546,313 hematopoietic cell states spanning these conditions. In addition to the MNC dataset introduced above, we included two progenitor-enriched datasets (*39*, *40*), a published atlas of hematopoiesis (*47*) and two *in vitro* datasets (*48*, *49*). While this heterogeneous corpus was used for representation learning, downstream biological analyses on *in vivo* cell states were restricted to internally generated datasets to minimize technical and annotation confounding.

The CD34^+^ atlas was downloaded from https://cellatlas.io/perturbgen. The Zeng dataset (*47*) was downloaded from cellxgene (https://cellxgene.cziscience.com/collections/f6c50495-3361-40ed-a819-fb9644396ed9) in December 2024. Stromal cells were excluded from subsequent analyses. The Li dataset (*39*) was downloaded from GEO (accession number GSE189161). The Sakurai dataset (*49*) was downloaded from GEO (accession number GSE192519). The Gao dataset (*48*) was downloaded from GEO (accession number GSE276896).

#### GP curation

We first defined a list of hematopoiesis transcription factors based on prior publications (*137–141*). We then obtained their target genes from CollecTRI (*50*). To maintain specificity while ensuring sufficient data, we retained only those transcription factors with a reported number of target genes between 25 and 250, and also removed regulons which had high overlap. Finally, we removed any GP that expressed fewer than 5 genes on average per cell in more than two datasets from the training corpus. We supplemented this list of transcription factors with signaling pathway response signatures from PROGENy (*31*), keeping 200 significant genes per GP as above. The final selection can be found in **Supplementary table t10**.

### Expansion cultures

#### Human Umbilical Cord Blood (hUCB) processing and CD34+ enrichment

CB samples were processed as previously described (*142*). Briefly, CB-derived mononuclear cells were isolated using density gradient centrifugation. Whole blood samples were first diluted 1:1 in PBS, then separated using Pancoll (PAN-Biotech) at 500 g for 25 minutes with brake off. Erythrocytes were lysed by incubation with ammonium chloride-based Red Blood Cell Lysis Buffer (BioLegend) for 15 minutes at 4°C in the dark. CD34+ selection was then performed using CD34 MicroBeads and FcR blocking reagent (both Miltenyi Biotec, 130-100-453) in PBS containing 3% fetal calf serum (FCS; PAN-Biotech, P40-37500) for 30 minutes at 4°C, followed by separation using LS Columns (Miltenyi Biotec, 130-042-401). Purified CD34+ cells were cryopreserved in FCS containing 10% DMSO (Sigma) and stored at -150°C until use.

#### 3a media and STEMSPAN cell cultures

CD34+ cells were seeded at 10,000 cells per well in CellBIND tissue culture plates (Fisher Scientific, 10510733). Cells were cultured at 37°C with 5% CO_2_ and 20% O_2_, in their corresponding media with varying concentrations, 1, 10 and 100 ng/mL, of SEC61 inhibitor (Cambridge bioscience, T40146-5mg) or DMSO (vehicle control). 3a media was composed of: IMDM medium (Gibco, 12440-053), 1% (v/v) Insulin-Transferrin-Selenium-Ethanolamine (ITS-X) (Gibco, 51500-056), 1% (v/v) Penicillin-Streptomycin-Glutamine (P/S/G) (Gibco, 10378-016), 0.1% Soluplus (BASF, 50539897), 1 μM 740Y-P (Selleckchem, S7865), 0.1 μM Butyzamide (Selleckchem, E1352) and 1 μM UM729 (MedChemExpress, HY-15972). StemSpan SFEM (Stem Cell Technologies, 09650) cultures either with 750 nMSR-1 (Selleckhem, S2858) or 35 nM UM171 (MedChemExpress, HY-12878) were supplemented with 100 ng/ml SCF (Qkine, Qk078-0100), 100 ng/mL rhTPO (Miltenyi, 130-094-013), 100 ng/mL FLT3L (Qkine, Qk087-0100) and 10 μg/mL human LDL (Stem Cell Technologies, 02698). Samples were cultured until day 9. Complete media changes were made every 2-3 days.

#### Spectral flow cytometry

Three technical replicates per CB were collected on day 9 of culture in V-bottom plates (Greiner Bio-one, 651261) and supernatant was removed after centrifugation at 300 g for 6 minutes at 4°C. Samples were resuspended with 50 μl of antibody cocktail mix, as described in (**Supplementary table t11**), and incubated on ice in the dark for 30 minutes. Subsequent to incubation, samples were washed once with PBS. Samples were then resuspended with 7AAD (1:1000) (Fisher Scientific, A1310). All cells were analysed by spectral flow cytometry using Sony ID 7000 Spectral Cell Analyzer and FCS files were analysed by Flowjo software 10.8.1. HSCs and GMPs were quantified as CD34^+^ CD45RA^−^ CD90^+^ EPCR^+^ and CD34^+^ CD7^−^ CD10^−^ CD45RA+ populations, respectively.

### Skin dataset

Datasets for scRNAseq and single-cell spatial transcriptomics were downloaded from https://cellatlas.io/studies/spatial-skin-atlas. Unpublished data analysed for this study, including validation sections for atopic dermatitis and the aligned TIFF files from spatial proteomics, will be made publicly available online upon publication at https://cellatlas.io/tripso.

#### Single cell RNA-seq data analysis

We downloaded the raw scRNA-seq data from the spatial skin atlas webportal. Gene expression was normalized to total counts per cell and log-transformed (log1p); for single-gene visualizations, values were min–max scaled per gene. Each GP score was computed from raw counts.

#### Spatial Transcriptomics analysis

We downloaded the Xenium spatial transcriptomics data from the spatial skin atlas webportal. Gene expression was library-size normalized (total-count normalization) and log-transformed (log1p). For single-gene visualizations, expression values were additionally min–max scaled per gene. Each GP score was computed from raw counts. For plotting gene expression in space, we plotted cell polygons using measured cell and nuclear area to preserve the true relative sizes of cells.

#### Quantitative assessment of spatial co-localization between gene programs and T cells hotspots

To quantify whether GP activity co-varied with the spatial organization of T cells, we computed the relationship between each cell’s GP module score and its distance to regions of high local density (“hotspots”) of a selected T-cell subtype on Xenium sections.

For each Xenium slide, we extracted 2D spatial coordinates for all segmented cells from the Xenium AnnData object. We then defined a target population of interest (e.g. T cells) and fit a two-dimensional Gaussian kernel density estimate (KDE) to the coordinates of target cells (Gaussian kernel; bandwidth set in pixel units). The KDE was evaluated on a regular grid spanning the full tissue coordinate range with a fixed grid step. To identify hotspots, we thresholded the KDE surface at a high percentile (e.g., the 92nd percentile) to obtain a binary hotspot mask and extracted connected components. Small components below a minimum area threshold were discarded to avoid spurious hotspots. For each remaining component, we computed a hotspot center using the center of mass of the thresholded region (mapped back to tissue coordinate units). If no hotspot passed filtering, the centroid of target-cell coordinates was used as a fallback hotspot center.

We then computed the Euclidean distance between each cell in the slide and the nearest hotspot center using a fast nearest-neighbor search (KD-tree). Distances were binned into fixed-width intervals (in pixel units), and within each distance bin we aggregated the GP module score of interest. For each bin we reported the mean module score and its standard error (SE) across cells, and visualized the dependence of GP activity on hotspot proximity as the mean profile with a mean ± SE ribbon. This analysis yields an interpretable quantitative readout of spatial co-dependence between GP activity and localized enrichments of a particular cell type on tissue sections.

#### Experimental validation of GP activity via Xenium spatial transcriptomics and Akoya Phenocycler spatial proteomics

In order to validate our observations of GP activity in AD and psoriasis, we leveraged an in-house dataset of AD and Psoriasis sections. We generated spatial transcriptomics for 8 novel skin sections, following the steps described in (*86*). Six-millimetre punch biopsies of adult human skin were obtained from patients diagnosed with moderate-to-severe AD or psoriasis. Diagnoses were confirmed by qualified dermatologists. Biopsies from AD and psoriasis patients were embedded in OCT compound and snap-frozen for downstream processing. Ethical approval for the collection and storage of all research material was granted by the relevant research ethics committees at St John’s Institute of Dermatology, Guy’s Hospital, London (REC reference: EC00/128 (AD samples) and 11/H0802/7 (Psoriasis samples)).

#### Xenium spatial transcriptomics

Fresh frozen human skin tissue (AD and psoriasis) embedded in OCT was sectioned at 10 µm thickness. Spatial transcriptomic profiling was performed using the 10x Genomics Xenium Prime 5K Human Pan-Tissue & Pathways panel following manufacturer’s instructions. Tissue sections were mounted within the Xenium slide capture area and processed according to the standard Xenium *in situ* gene expression workflow, including antibody-based cell segmentation. Following imaging, slides were maintained in PBS at 4 °C for downstream histological and protein-based analyses, including H&E staining and multiplex immunofluorescence. Patient demographic characteristics and Xenium QC metrics can be found in **Supplementary table t12**.

#### Multiplex immunofluorescence (Akoya PhenoCycler)

After the Xenium assay, slides underwent multiplex immunofluorescence using the Akoya PhenoCycler-Fusion platform with the Human IO60-plex panel. Staining was performed according to the manufacturer’s one-step protocol, omitting additional acetone and paraformaldehyde pre-fixation steps. The PhenoCycler-Fusion system performs iterative cycles of reporter hybridization, imaging, and reporter removal via isothermal washes until all targets are acquired. Whole-slide scans were generated upon completion of the assay. Image processing and visualization of marker expression and co-expression were conducted using QuPath, and exported images were saved in TIFF format.

#### Spatial Proteomics analysis

We analyzed registered multi-channel CODEX OME-TIFF images with matched Xenium data. To align the mosaic QPTIFF image, we used *palom* (*143*). For each slide, we matched the aligned OME-TIFF to the Xenium segmentation output by slide identifier, and processed each slide independently.

Xenium cell boundaries were provided as polygon vertex coordinates in microns in *cell_boundaries.csv.gz* inside each Xenium bundle. Vertex coordinates were converted to pixel indices using the Xenium pixel size (p=0.2125 μm per pixel). Polygons were rasterized into a 2D label image. In parallel, we stored a one-to-one mapping between cell IDs and each polygon, which was later attached to the quantified proteomics table for cross-modality linkage. For each slide, we constructed a *SpatialData* object containing (i) the aligned CODEX image as a 3D array with dimensions (*c*, *y*, *x*) and channel names provided in the experiment channel list, and (ii) the rasterized Xenium segmentation label image with dimensions (*y*, *x*). This ensured that per-cell protein quantification was performed in the same pixel reference frame as Xenium segmentation.

To reduce imaging noise and stabilize downstream quantification, we applied channel-wise intensity processing using the *spatialproteomics* package (*144*). Specifically, we performed per-channel quantile-based thresholding (quantile *q* = 0.95), followed by a 2D median filter (kernel size *k* = 3) applied to each channel. Protein expression was quantified per Xenium-segmented cell by computing the mean pixel intensity within each cell mask for each channel. Quantified intensities were then transformed using an *arcsinh* transform to compress dynamic range. Z-score transformation was computed for each channel. For each slide, per-cell protein measurements were exported and integrated with the corresponding Xenium AnnData object, enabling downstream spatial protein visualization and direct comparison to Xenium-derived transcriptomic signals (e.g., CD45/PTPRC).

