## Supplementary Figures for "Self-supervised learning for a gene program-centric view of cell states"

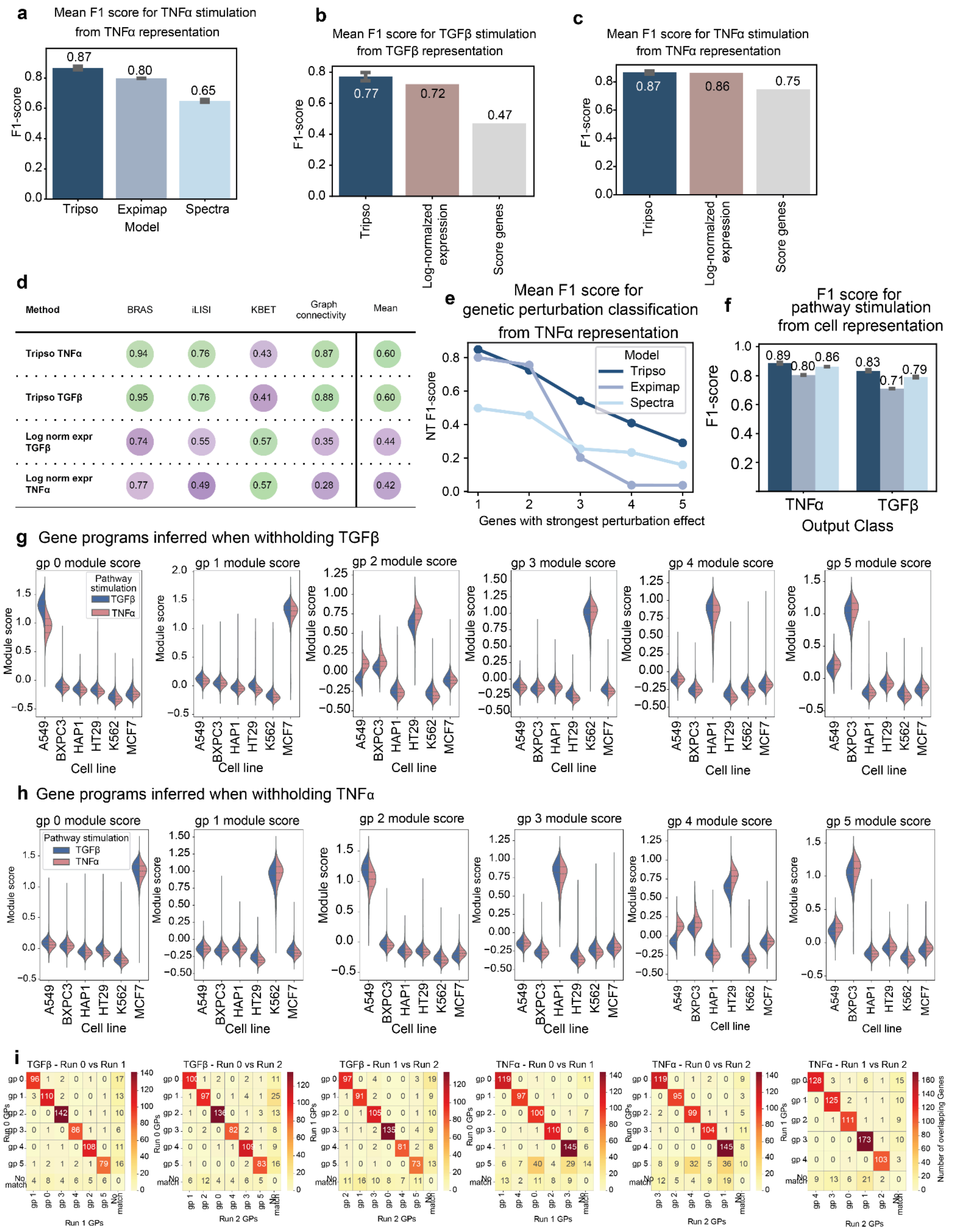


**Supplementary Figure S1: Benchmarking Tripso** (**a**) F1-score for TNFα classification from TNFα activity representation learned by each model. (**b/c**) F1-score for TGFβ (b) TNFα (c) classification based on the GP representation learned by Tripso or simple linear baselines. (**d**) Batch integration metrics for GP representations learned by Tripso or obtained by concatenated log normalized expression of GP genes (higher is better, more purple =

lower, more green = higher). (**e**) F1-score for classifying non-targeting controls from TNFα activity representation learned by each model. The x-axis represents the number of perturbed gene classes; genes were selected based on the strength of the induced perturbation. (**f**) F1-score for classifying pathway stimulation score based on the global cell representation learned by each model. (**g/h**) Gene expression module score for each of the GPs inferred by Tripso. Values on the x-axis represent different cell lines, split by cytokine stimulation. Each subplot corresponds to one GP. (**g**) GP inferred in the configuration where Tripso is trained with only TNFα as known GP, (**h**) GP inferred in the configuration where Tripso is trained with only TGFβ as known GP. (**i**) Evaluation of the overlap between Tripso-inferred GPs using different random seeds. The first three panels correspond to the configuration where Tripso is trained with only TNFα as known GP, the last three panels correspond to the configuration where Tripso is trained with only TGFβ as known GP. Within each panel, each row/column corresponds to a GP inferred by Tripso, where the GP on the rows all come from a run with the same random seed, and the GP on the columns all come from a run with the same configuration but a different random seed.

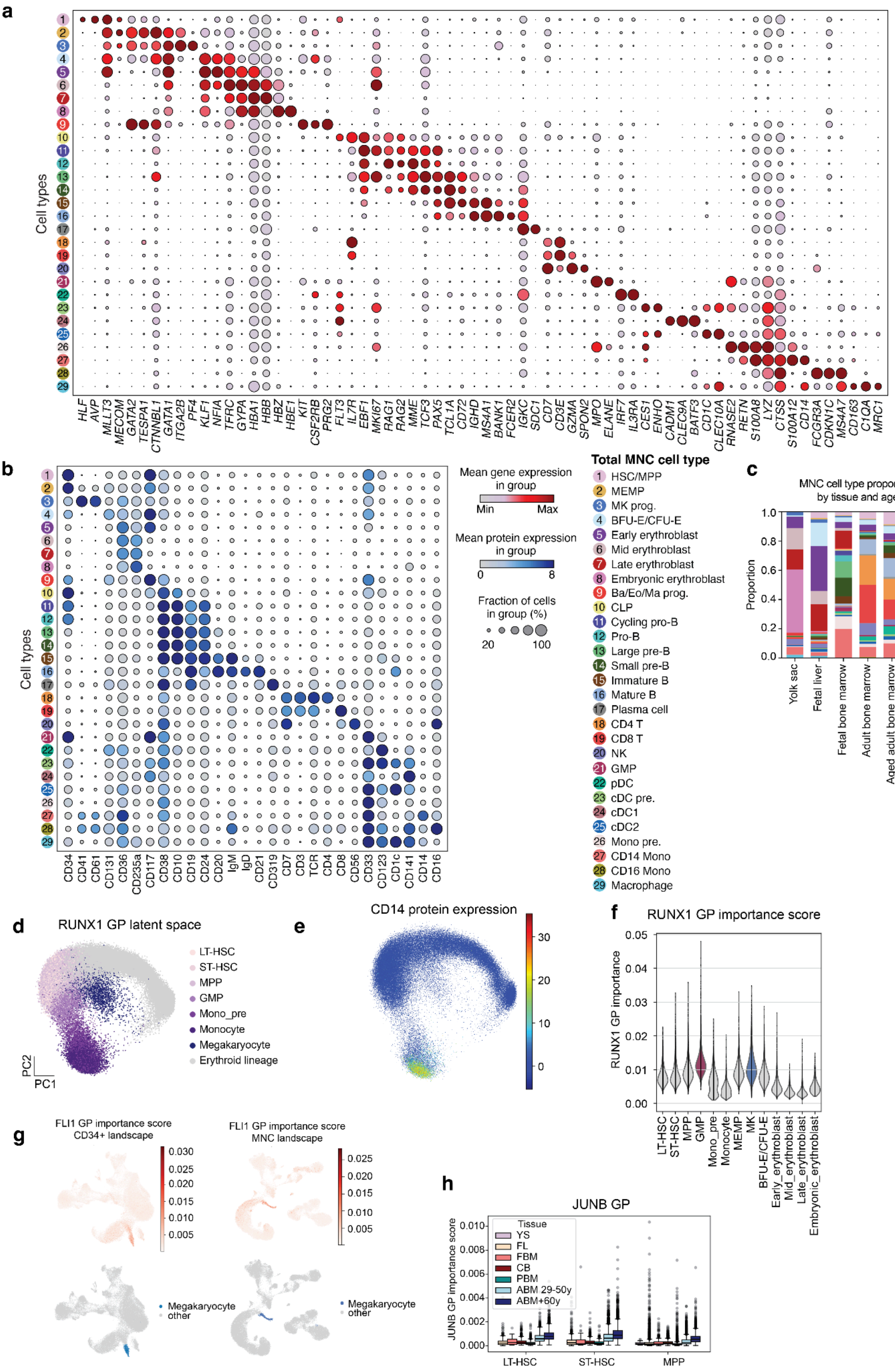
a
Cell types
HLF AVP MLLT3 MECOM GATA2 TESPA1 CTNNBL1 GATA1 ITGA2B PF4 KLF1 NFIA TFRC GYPA HBA1 HBB HBZ HBE1 KIT CSF2RB PRG2 FLT3 IL7R EBF1 MKI67 RAG1 RAG2 MME TCF3 PAX5 TCL1A CD72 IGHD MS4A1 BANK1 FCER2 IGKC SDC1 CD7 CD3E GZMA SPON2 MPO ELANE IRF7 IL3RA CES1 ENHO CADM1 CLEC9A BATF3 CD1C CLEC10A RNASE2 RETN S100A8 LYZ CTSS S100A12 CD14 FCGR3A CDKN1C MS4A7 CD163 C1QA MRC1
b
Cell types
CD34 CD41 CD61 CD131 CD36 CD235a CD117 CD38 CD10 CD19 CD24 CD20 IgM IgD CD21 CD319 CD7 CD3 TCR CD4 CD8 CD56 CD33 CD123 CD1c CD141 CD14 CD16
Mean gene expression in group
Min
Max
Mean protein expression in group
0
8
Fraction of cells in group (%)
20
100
Total MNC cell type
1 HSC/MPP
2 MEMP
3 MK prog.
4 BFU-E/CFU-E
5 Early erythroblast
6 Mid erythroblast
7 Late erythroblast
8 Embryonic erythroblast
9 Ba/Eo/Ma prog.
10 CLP
11 Cycling pro-B
12 Pro-B
13 Large pre-B
14 Small pre-B
15 Immature B
16 Mature B
17 Plasma cell
18 CD4 T
19 CD8 T
20 NK
21 GMP
22 pDC
23 cDC pre.
24 cDC1
25 cDC2
26 Mono pre.
27 CD14 Mono
28 CD16 Mono
29 Macrophage
c
MNC cell type proportion by tissue and age
Proportion
1.0
0.8
0.6
0.4
0.2
0.0
Yolk sac
Fetal liver
Fetal bone marrow
Adult bone marrow
Aged adult bone marrow
Cord blood
d
RUNX1 GP latent space
LT-HSC
ST-HSC
MPP
GMP
Mono_pre
Monocyte
Megakaryocyte
Erythroid lineage
PC2
PC1
e
CD14 protein expression
30
20
10
0
f
RUNX1 GP importance score
RUNX1 GP importance
0.05
0.04
0.03
0.02
0.01
0.00
LT-HSC
ST-HSC
MPP
GMP
Mono_pre
Monocyte
MEMP
MK
BFU-E/CFU-E
Early_erythroblast
Mid_erythroblast
Late_erythroblast
Embryonic_erythroblast
g
FLI1 GP importance score CD34+ landscape
0.030
0.025
0.020
0.015
0.010
0.005
FLI1 GP importance score MNC landscape
0.025
0.020
0.015
0.010
0.005
Megakaryocyte
other
Megakaryocyte
other
h
JUNB GP
JUNB GP importance score
0.010
0.008
0.006
0.004
0.002
0.000
Tissue
YS
FL
FBM
CB
PBM
ABM 29-50y
ABM+60y
LT-HSC
ST-HSC
MPP

**Supplementary Figure S2: Tripso identifies age-specific cell states in hematopoiesis**: (**a**) Dotplot showing key markers used to annotate each cell type in CITE-seq. Color intensity reflects min-max normalized gene expression of each gene. (**b**) Dotplot showing key markers used to annotate each cell type in CITE-seq dataset. Color intensity reflects min-max normalized gene expression of each surface marker. (**c**) Proportion of cell types by tissue, cell type colors as in a/b. (**d/e/f**) Scatter plot on the first two principal components of stem, erythroid, monocytic and megakaryocyte cells in the RUNX1 GP latent space, colored by d) cell type, e) CD14 surface protein expression (CITE-seq), f) RUNX1 GP importance score (GMP and megakaryocytes, which have the highest importance scores, are highlighted). (**g**) FLI1 GP importance score (learned by Tripso) projected onto the original gene expression UMAP (CD34+- left and MNC- right). Color intensity reflects FLI1 importance score, highlighted cell types are megakaryocytes (shown below). (**h**) JUNB GP importance score for LT-HSC, ST-HSC and MPP, grouped by tissue.

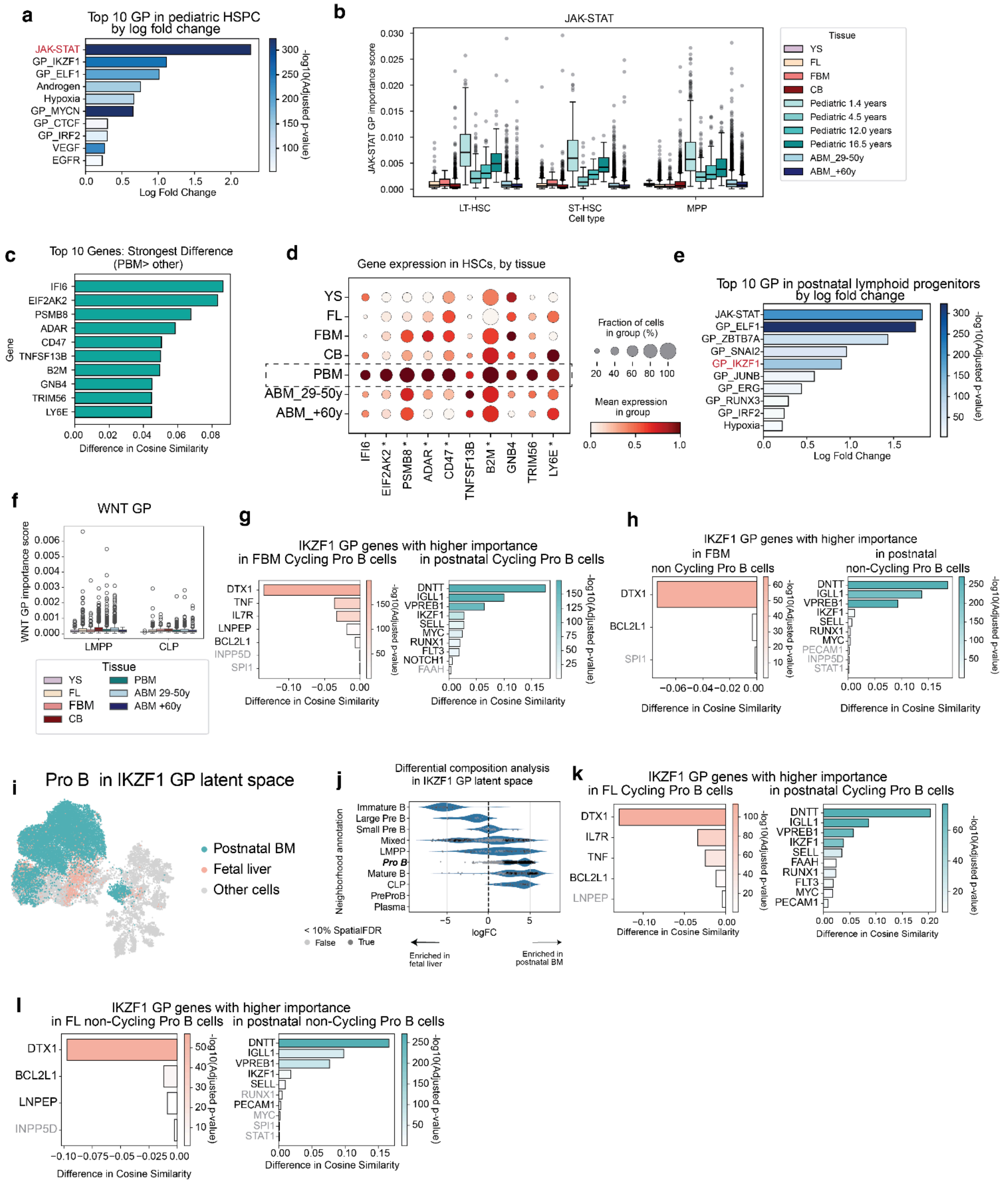

**Supplementary Figure S3: Tripso identifies age-specific cell states in hematopoiesis**
(**a**) Differential GP importance scores in pediatric HSC (LT-HSC, ST-HSC, MPP) compared to all other tissues, GP are ranked by log-fold change and colored by statistical significance. JAK-STAT (shown in Main Figure 3) is highlighted in red. (**b**) JAK-STAT GP importance

scores in HSC populations, separated by tissue. For pediatric donors, each bar represents one donor (ordered by age). (**c**) Differential gene importance for JAK-STAT GP in HSCs in pediatric bone marrow compared to other tissues. Adjusted p-values $<10^{-12}$. (**d**) Dotplot showing min-max normalized expression of the top 10 JAK-STAT GP genes showing differential importance in pediatric bone marrow. * indicates genes which belong to MsigDB Hallmarks Interferon Alpha response. (**e**) Differential GP importance scores in postnatal lymphoid multipotent progenitors (LMPP) and common lymphoid progenitors (CLP) compared prenatal LMPP and CLP, GP are ranked by log-fold change and colored by statistical significance. IKZF1 (shown in Main Text Figure 3) is highlighted in red. (**f**) WNT GP importance score for LMPP and CLP, grouped by tissue. (**g/h**) Differential gene importance for IKZF1 GP in cycling **(g**) and non-cycling (**h**) pro-B cells in postnatal bone marrow (ABM, PBM) compared to fetal bone marrow (FBM). (**i**) UMAP of B lineage cells from fetal liver or post natal bone marrow in IKZF1 GP embedding space, pro-B cells are highlighted and colored by tissue stage. (**j**) Milo differential composition analysis results, testing for neighborhoods enriched in postnatal bone marrow (PBM, ABM) vs fetal liver. Darker circles represent spatial FDR < 10%. (**k/l**) Differential gene importance for IKZF1 GP in cycling (k) or non-cycling (l) pro-B cells in postnatal bone marrow (ABM, PBM) compared to fetal liver. In panels g/h/k/l Gene names in gray correspond to differences with adjusted p-values >0.05. All differences in cosine similarity scores represent averages across 3 model training runs.

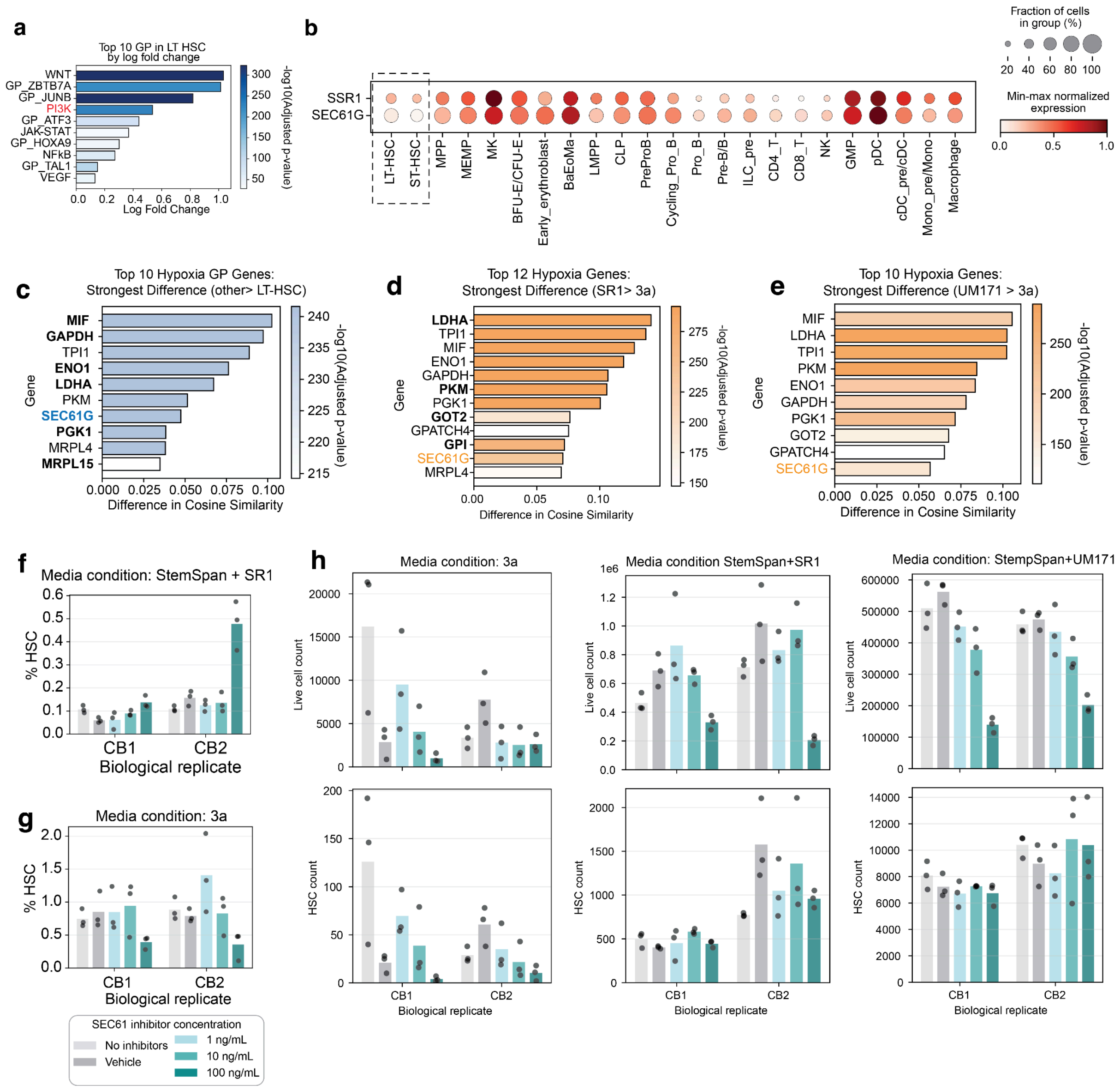


**Supplementary Figure S4: Gene program-guided *in vivo*–*in vitro* mapping and perturbation prioritization in hematopoiesis:** (**a**) Differential GP importance scores in transcriptomically defined LT-HSC compared to stem and progenitor cells. GP are ranked by log-fold change and colored by statistical significance. PI3K (shown in Main Figure 4) is highlighted in red. (**b**) Log normalized expression of ER translocon components SSR1 and SEC61G *in vivo* (CD34+ dataset, all tissues and stages). (**c**) Differential gene importance for Hypoxia GP genes in stem and progenitor cells compared to LT-HSC *in vivo.* (**d**) Differential gene importance for Hypoxia GP genes in HSPCs cultured in SR-1 media compared to 3a media. (**e**) Differential gene importance for Hypoxia GP genes in HSPCs cultured in UM171 media compared to 3a media. All GP importance scores and gene cosine similarity values represent averages across 3 random seeds. Gene names in bold correspond to genes which were significantly differently expressed (adjusted p-value <0.05) in pseudobulked differential gene expression. (**f/g**) Proportion of phenotypic HSCs detected under different concentrations of SEC61 inhibitor or control (no inhibitors and vehicle) in SR-1 (f) or 3a (g) media). (**h**) Total number of live cells (top row) or immunophenotypic HSCs (bottom row) in 3a media (left column) SR-1 (middle) or UM171 (right) conditions with different concentrations of SEC1 inhibitor. Bar colors represent concentration as shown in (f/g). Each cord blood (CB) represents a biological replicate, and each dot represents a technical replicate.

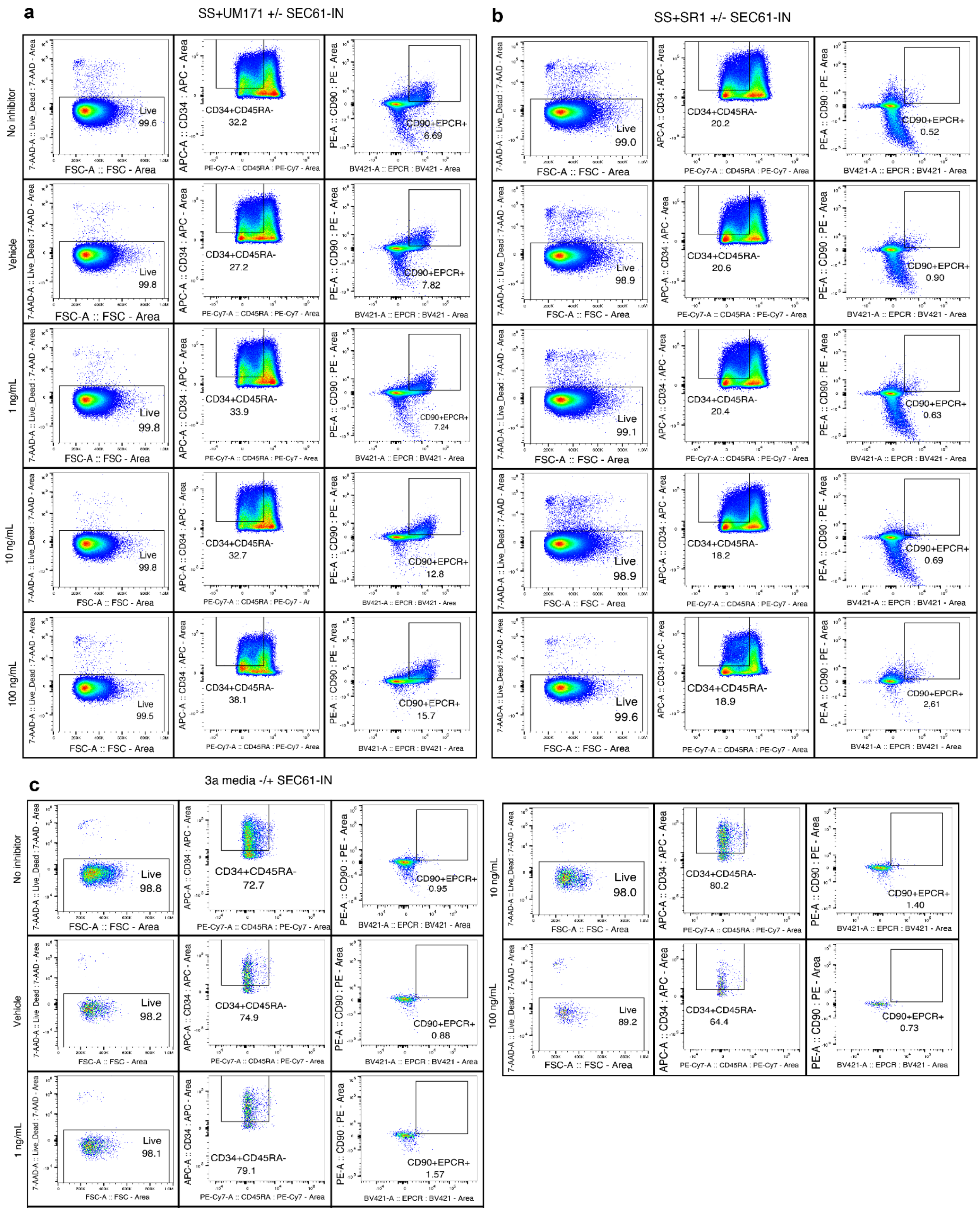


**Supplementary Figure S5 : Gating strategy for expansion culture experiments.** (**a**) StemSpan (SS) with UM171, (**b**) StemSpan (SS) with SR-1, and (**c**) 3a media conditions. Rows indicate SEC61 inhibitor conditions (Control, Vehicle, 1 ng/mL, 10 ng/mL, 100 ng/mL).

Within each condition, panels show sequential gating of live cells, $CD34^{+}CD45RA^{-}$ cells, and $CD90^{+}EPCR^{+}$ cells.

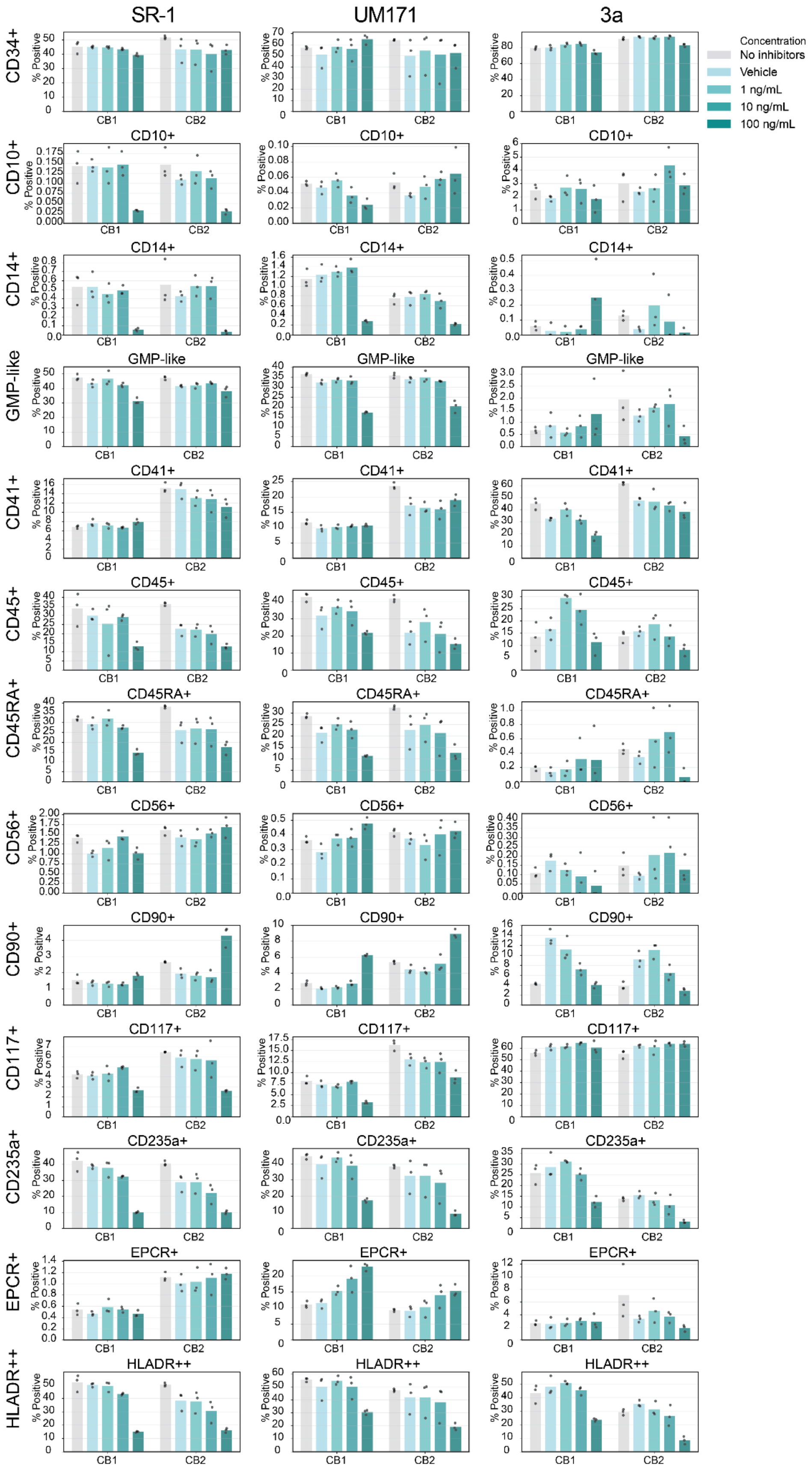


**Supplementary Figure S6**: Flow cytometry data showing the proportion of live cells assigned to each surface marker population. Each row corresponds to a marker indicated on the left-hand side. Each column corresponds to a cell culture condition (SS + UM171, SS + SR1, 3a). Bar colors represent SEC61 treatment concentration. Each cord blood (CB) represents a biological replicate, and each dot represents a technical replicate. GMP-like cells are defined as CD34+ CD7− CD10− CD45RA+.

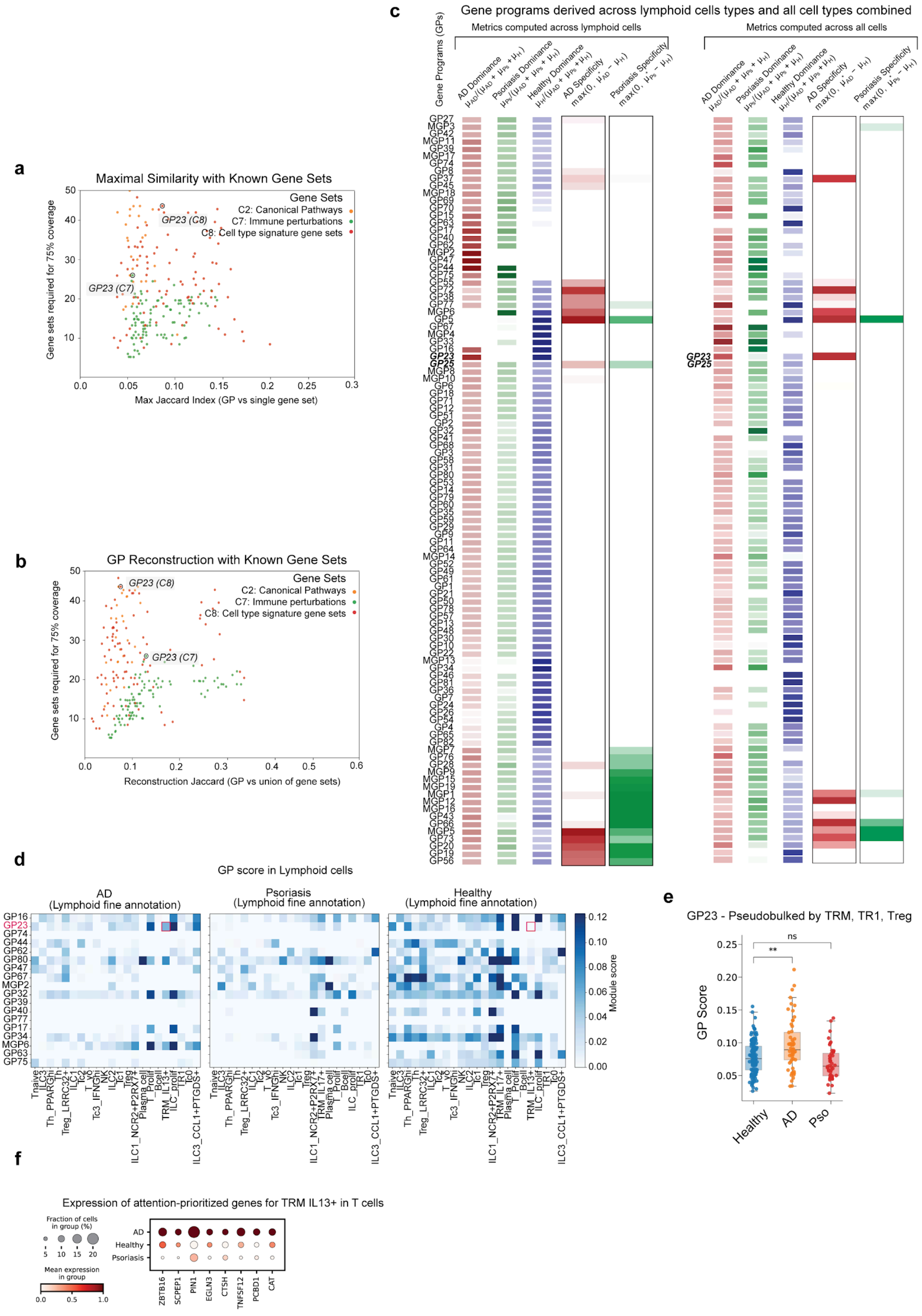


**Supplementary Figure S7 Prioritization and characterization of Tripso-derived GPs in lymphoid cells:** (**a**) Best single-set similarity during GP reconstruction. For each

Tripso-derived GP and each curated collection (Hallmarks H, Canonical Pathways C2, Immunologic Signatures C7, and Cell Type Signatures C8), we computed the maximum Jaccard similarity between the GP and its best-matching gene set in that collection (x axis). The y axis reports the number of gene sets selected by the false positive–penalized greedy reconstruction procedure for that GP-collection pair. Each dot corresponds to one GP-collection pair (thus each GP appears once per collection) and is colored by collection. (**b**) GP reconstructability from curated resources. For each GP and collection, we attempted to reconstruct the GP using a false positive–penalized greedy selection of gene sets until reaching the target coverage threshold (≥75% of GP genes) or a maximum set limit (Methods). Reconstruction quality is summarized by the Jaccard similarity between the GP and the union of selected gene sets (x axis) and by the number of gene sets required to reach the coverage threshold (y axis). Points exceeding the displayed y-axis range are omitted. Each dot corresponds to one GP-collection pair and is colored by collection. (**c**) Condition-level prioritization of GPs by dominance and specificity. Heatmaps show GP dominance across AD, psoriasis, healthy skin and disease specificity relative to healthy (AD−Healthy, psoriasis−Healthy; Methods), computed either within lymphoid cells (left) or across all cells (right). GP23 and GP25 are highlighted. Values are displayed with linear white-to-color scales (red for AD, green for psoriasis, blue for healthy). (**d**) Lymphoid-focused prioritization across diseases. Heatmaps show mean module scores for selected lymphoid-associated GPs (rows), prioritized by high dominance or specificity in AD or psoriasis, across fine lymphoid subsets (columns) for AD, psoriasis, and healthy skin. GP23 is highlighted in the IL13+ TRM subset (absent in psoriasis). (**e**) Pseudobulk comparison of GP23 across regulatory T cell subsets. GP23 pseudobulk module scores for TRM, TR1, and Treg subsets, aggregated per patient and stratified by condition. Each dot represents one patient; statistical testing uses Welch’s t test. GP23 differs between AD and healthy ($p < 0.001$), whereas psoriasis versus healthy is not significant (ns). (**f**) Expression of GP23 constituent genes prioritized by differential analysis of Tripso attention scores in $IL13^{+}$ TRM cells relative to other T cells. Dot plot showing expression patterns of a subset of GP23 genes across T cells in AD, healthy skin, and psoriasis. Dot color indicates mean expression (scaled 0–1 per gene across conditions), and dot size indicates the fraction of T cells expressing each gene.

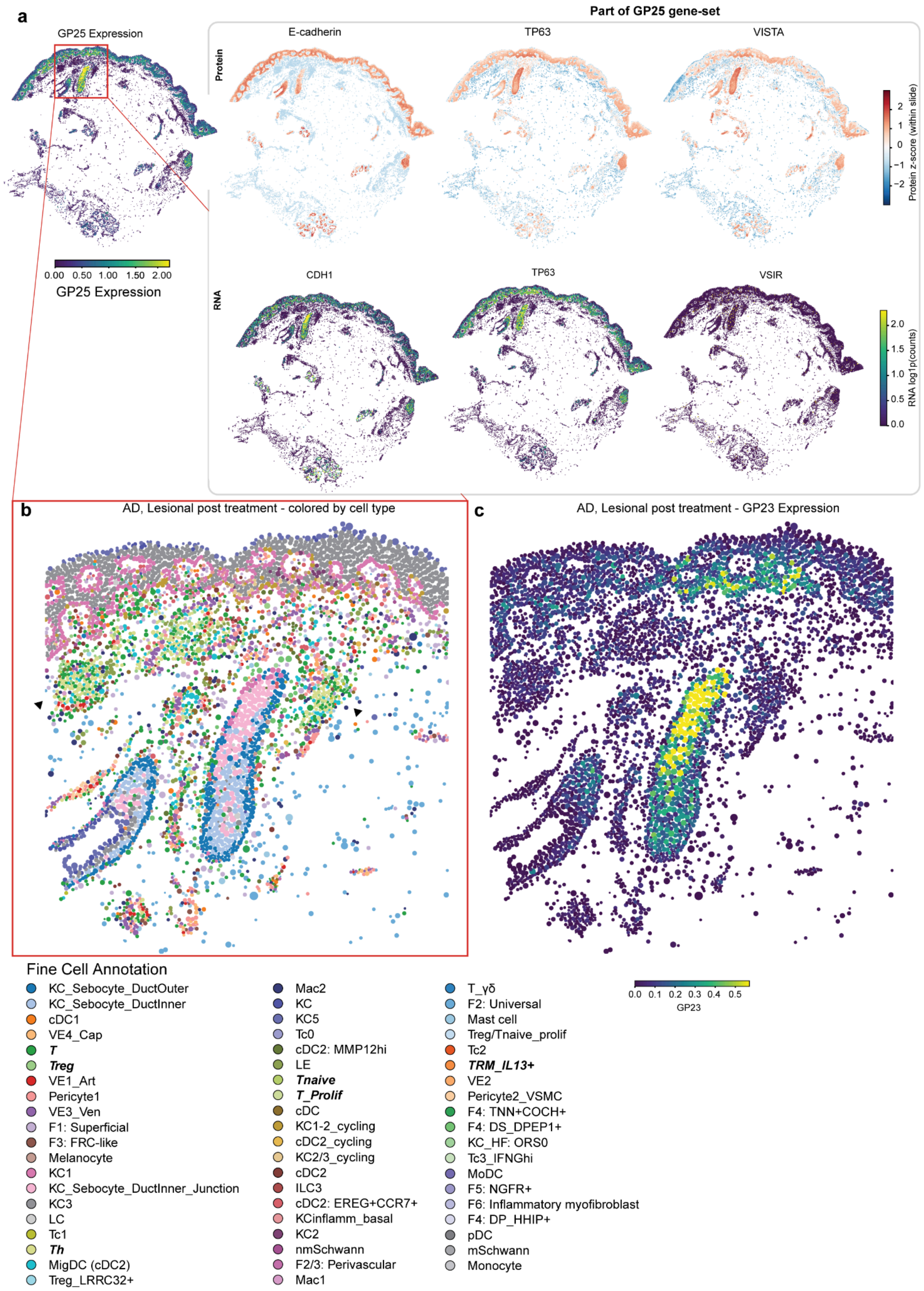


**Supplementary Figure S8: Spatial proteomics support for GP25 markers and GP23 enrichment near T cell aggregates: (a)** Orthogonal validation of GP25 constituent markers by matched spatial proteomics and spatial transcriptomics in an AD section. Left, spatial distribution of GP25 program activity (module score). Top row, spatial proteomics signal for

E-cadherin, TP63, and VISTA, values expressed as Z-score enrichment for the protein signal (Methods). Bottom row, corresponding Xenium RNA signals for their encoding genes (CDH1, TP63, and VSIR), shown in the same tissue coordinate frame. Values plotted in log1p. **(b)** Fine cell-type context in a $CD45^{+}$, sebaceous-associated immune niche. Zoomed region with fine cell-type annotations; a cluster of T helper cells is indicated (black triangles). Legends report the cell types for the same region. In shades of color green: T, Th, T naive, T proliferant. **(c)** Spatial expression of GP23 in the same zoomed region as (b), showing a strong GP23 signal adjacent to the T helper cell cluster near the sebaceous gland/ductal area.

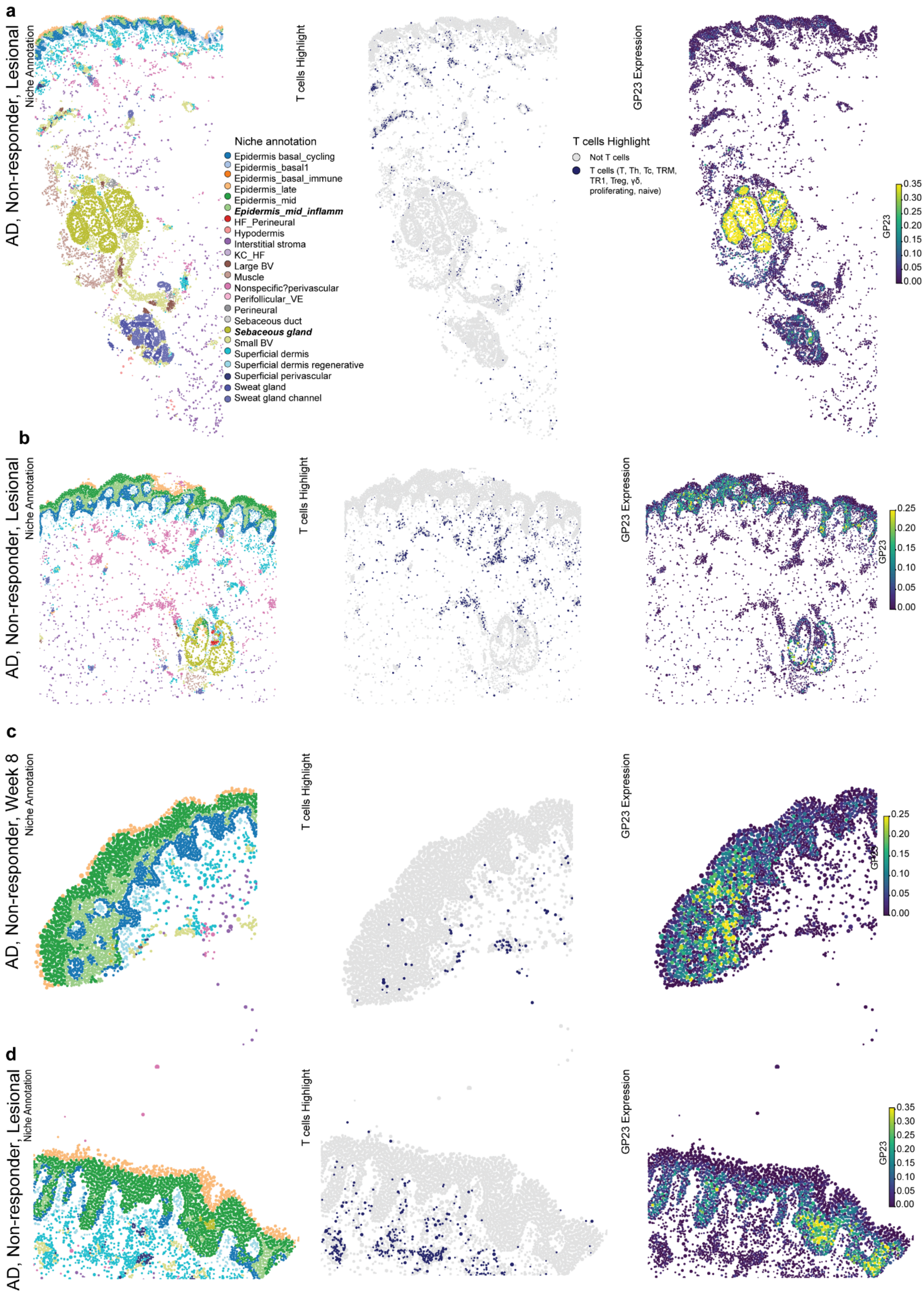


**Supplementary Figure S9: GP23 spatial validation in independent Xenium sections from AD lesional skin: (a–d)** Zoomed regions from four in-house Xenium sections from atopic dermatitis non-responder patients, selected to visualize GP23-positive areas. For each section, the left panel shows spatial niche annotations, the middle panel highlights T cells within the same region, and the right panel shows GP23 program activity (module score) across cells. Niche and T cell legends are shared across panels; GP23 is displayed on a continuous scale (higher values indicate higher program activity).

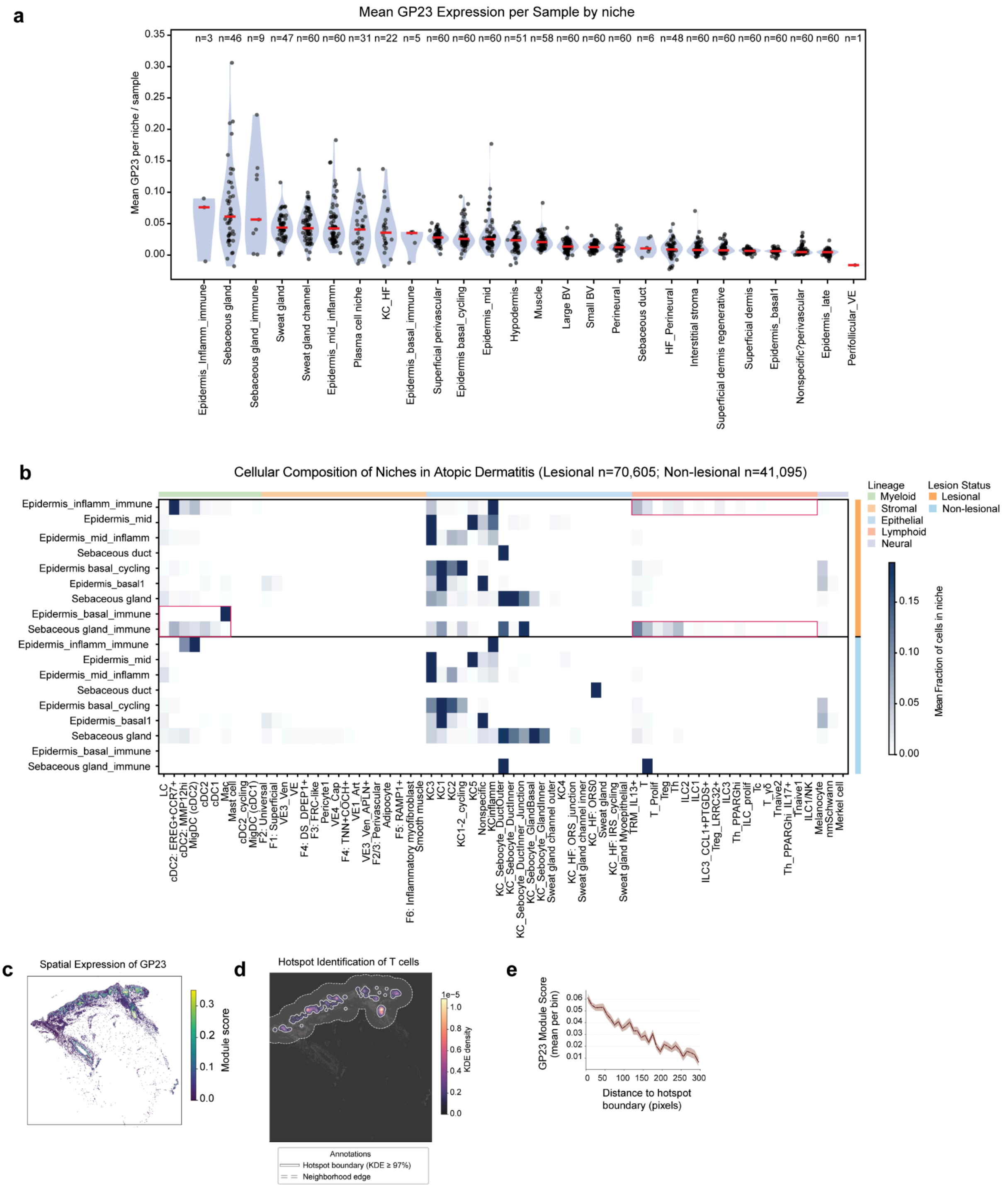


**Supplementary Figure S10: GP23 niche enrichment and spatial proximity to T cell hotspots:** (**a**) GP23 program activity across spatial niches. Violin plots show the distribution of GP23 module scores across cells within each annotated spatial niche. Each dot represents the per-sample mean GP23 score within that niche; niches are ordered by decreasing mean GP23 activity. (**b**) Mean niche cellular composition by lesion status. Heatmaps show the mean fraction of annotated cell types (columns, ordered by lineage) within each spatial niche (rows), stratified by lesion status. Each tile represents the average cell-type composition for a given niche-cell type pair within the corresponding lesion-status group. Red boxes highlight niches (*Sebaceous gland_immune* and *Inflammatory epidermis_immune*) showing increased myeloid and lymphoid representation in lesional tissue. (**c**) Spatial distribution of GP23 in a resolved psoriasis section. Cell-level GP23 module scores are shown on the tissue section. (**d**) Identification of T cell hotspots in the same section as (c). T cell local density was estimated by kernel density estimation (KDE; Methods) and visualized with a magma scale. Hotspot boundaries were defined as KDE ≥ 97th percentile; the neighborhood edge denotes a 400-pixel offset from the hotspot boundary. (**e**)  GP23 activity as a function of distance to T cell hotspots. The x axis shows distance (pixels) from the nearest hotspot boundary defined in (d), and the y axis shows the mean GP23 module score per distance bin (line; ± standard error of the mean).

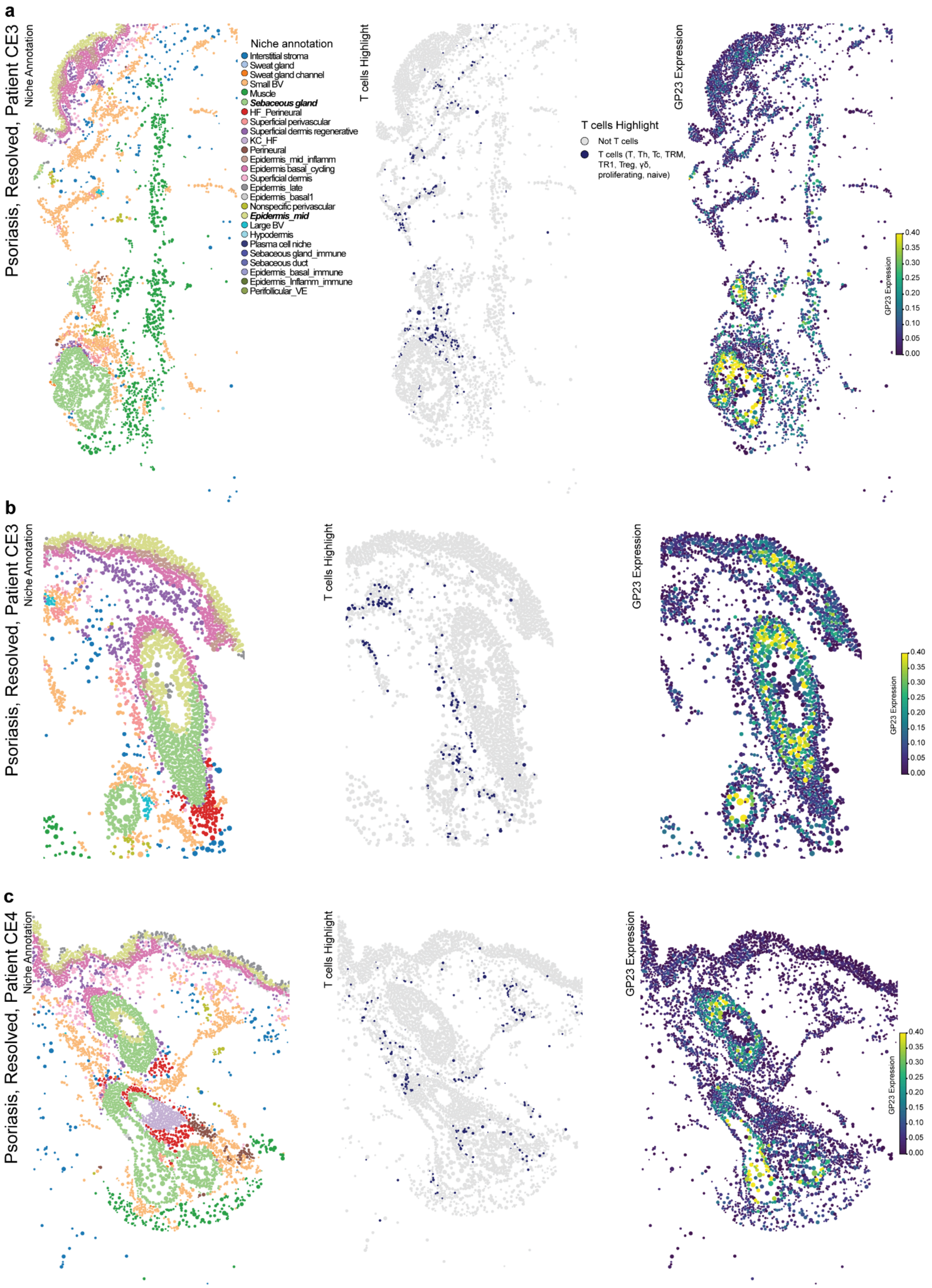


**Supplementary Figure S11: GP23 spatial patterns in resolved psoriasis sections: (a-c)** Zoomed regions from resolved (lesion-cleared) psoriasis Xenium sections. Panels a and b show two regions from the same patient, whereas c is from a different patient. For each region, the left panel shows spatial niche annotations, the middle panel highlights T cells, and the right panel shows GP23 program activity (module score) across cells. Niche and T cell legends are shared across panels; GP23 is displayed on a continuous scale (higher values indicate higher program activity).
